# Social recognition in rats and mice requires integration of olfactory, somatosensory and auditory cues

**DOI:** 10.1101/2020.05.05.078139

**Authors:** Shani Haskal de la Zerda, Shai Netser, Hen Magalnik, Mayan Briller, Dan Marzan, Sigal Glatt, Shlomo Wagner

## Abstract

In humans, discrimination between individuals, also termed social recognition, can rely on a single sensory modality, such as vision. By analogy, social recognition in rodents is thought to be based upon olfaction. Here, we hypothesized that social recognition in rodents relies upon integration of olfactory, auditory and somatosensory cues, hence requiring active behavior of social stimuli. Using distinct social recognition tests, we demonstrated that adult male rats and mice do not recognize familiar stimuli or learn the identity of novel stimuli that are inactive due to anesthesia. We further revealed that impairing the olfactory, somatosensory or auditory systems prevents recognition of familiar stimuli. Finally, we found that familiar and novel stimuli generate distinct movement patterns during social discrimination and that subjects react differentially to the movement of these stimuli. Thus, unlike what occurs in humans, social recognition in rats and mice relies on integration of information from several sensory modalities.

## Introduction

The ability to recognize or discriminate between individual conspecifics is crucial for the survival of members of gregarious species, as such ability guides appropriate interactions of these individuals with their social environment ^1-4^. In the literature, social recognition is used as a generic term for both the ability of a subject to categorize conspecifics into different classes, such as sex, genetic relatedness and familiarity (henceforth termed social recognition), as well as for the ability to recall the learned idiosyncratic identity of a specific individual based on a previous encounter, also termed individual recognition ^5,6^. In humans, social recognition can be based on cues detected by single sensory modalities. For example, humans can recognize a familiar person just by looking at their face (visual modality) or hearing their voice (auditory modality) ^7-9^. Moreover, human social recognition can occur even without active engagement with a social partner, such as by looking at a sleeping individual. Such single-modality based social recognition appears to also hold true for other primates ^10^. The generality of this ability among mammals remains, however, unclear. Mice and rats, the main mammalian laboratory models used in biomedical research ^11^, are social species known to exhibit social recognition ^12-17^. Specifically, during social interactions, these animals display higher investigative behavior towards novel conspecific individuals (henceforth termed social stimuli), as compared to those with whom they are familiar ^18^. Thus, in a social discrimination test, shorter times are dedicated by subjects for investigating a familiar stimulus, as compared to a novel one, reflecting recognition of the familiar stimulus. This type of social recognition, which is frequently used in the field of social neuroscience to assess typical social behavior ^19-22^, is widely assumed to be mediated by chemosensory cues released by the stimulus and received by the main and accessory olfactory systems of the subject ^13,14,17,23-26^. Therefore, in analogy to the human face, the identity of a social stimulus is thought to be represented by the passive signature of chemosensory cues (i.e. the olfactory signature), which distinguishes conspecifics ^14,27^. Still, despite reports that social recognition is impaired in anosmic animals ^28-30^, the reliance of social recognition in mice and rats solely on chemosensory cues has yet to be proven. Moreover, recent work showed that rat hippocampal CA1 pyramidal neurons respond to social cues, triggering the somatosensory and auditory systems in a social stimulus-specific manner ^31^ and that touch may by a crucial component of the social reward associated with social place preference ^32^. These studies raise the possibility that mice and rats integrate multimodal information during social interactions which can serve as a more complex basis for social recognition than the olfactory signature alone ^33^.

Accordingly, we challenged the common assumption that social recognition in mice and rats is solely based upon chemosensory cues in the present study. We instead hypothesized that social recognition relies upon the integration of olfactory, auditory and somatosensory cues, hence requiring active behavior of social stimuli.

## Results

### Mice and rats do not discriminate between anesthetized novel and familiar social stimuli

To examine possible involvement of stimulus-related behavior in rodent social recognition, we employed our published behavioral system (Netser et al. 2017), which allows automated and precise detection of bouts of investigative behavior towards social and non-social stimuli (Supp. Fig. 1A). Using this system, we first analyzed the time dedicated by C57BL/6J mice to investigate a cage-mate (CM) and a novel social stimulus (Novel), each located in triangular chambers at opposite corners of the experimental arena (familiarity discrimination test; Fig. 1A). The subject mice exhibited a clear preference for the novel social stimulus over the CM when these stimuli were awake, as reflected by the significantly longer time the subjects investigated the novel stimulus (Stimulus x Group: F(1,58)=7.406, p=0.005, mixed model ANOVA; *post hoc*: t(31)=5.622, p<0.001 paired t-test; Fig. 1C). We then exploited the unique design of our system to spread anesthetized social stimuli over the metal mesh of the triangular chamber (Supp. Fig. 1A) such that the ventral side of the anesthetized animal, including facial and anogenital regions, were accessible for investigation by the subject. We reasoned that if social recognition is solely based upon chemosensory signatures passively transmitted by social stimuli, social recognition should also transpire even with anesthetized stimuli. However, if active behavior of the stimulus is also required for social recognition, then subjects would not be able to distinguish between anesthetized stimuli. We found that mice subjects completely lost their preference for the novel stimulus, relative to the CM, when both stimuli were anesthetized (*post hoc*: t(27)=0.969, p=0.171, paired t-test; Fig. 1C). Importantly, the total time of investigation slightly increased when the stimuli were anesthetized as compared to awake individuals (t(58)=-2.479, p=0.008, independent t-test; Fig. 1D), suggesting that the subjects were interested in the anesthetized animals at least as much as they were in awake stimuli. Similar results were found for Sprague-Dawley (SD) rats using the same paradigm (Stimulus x Group: F(1,29)=12.516, p<0.001, mixed model ANOVA; Supp. Fig. 2). These results suggest that social recognition in both mice and rats relies on the behavior of the social stimuli, such that subjects do not distinguish between novel and familiar stimuli if these stimuli are anesthetized.

**Figure 1.**
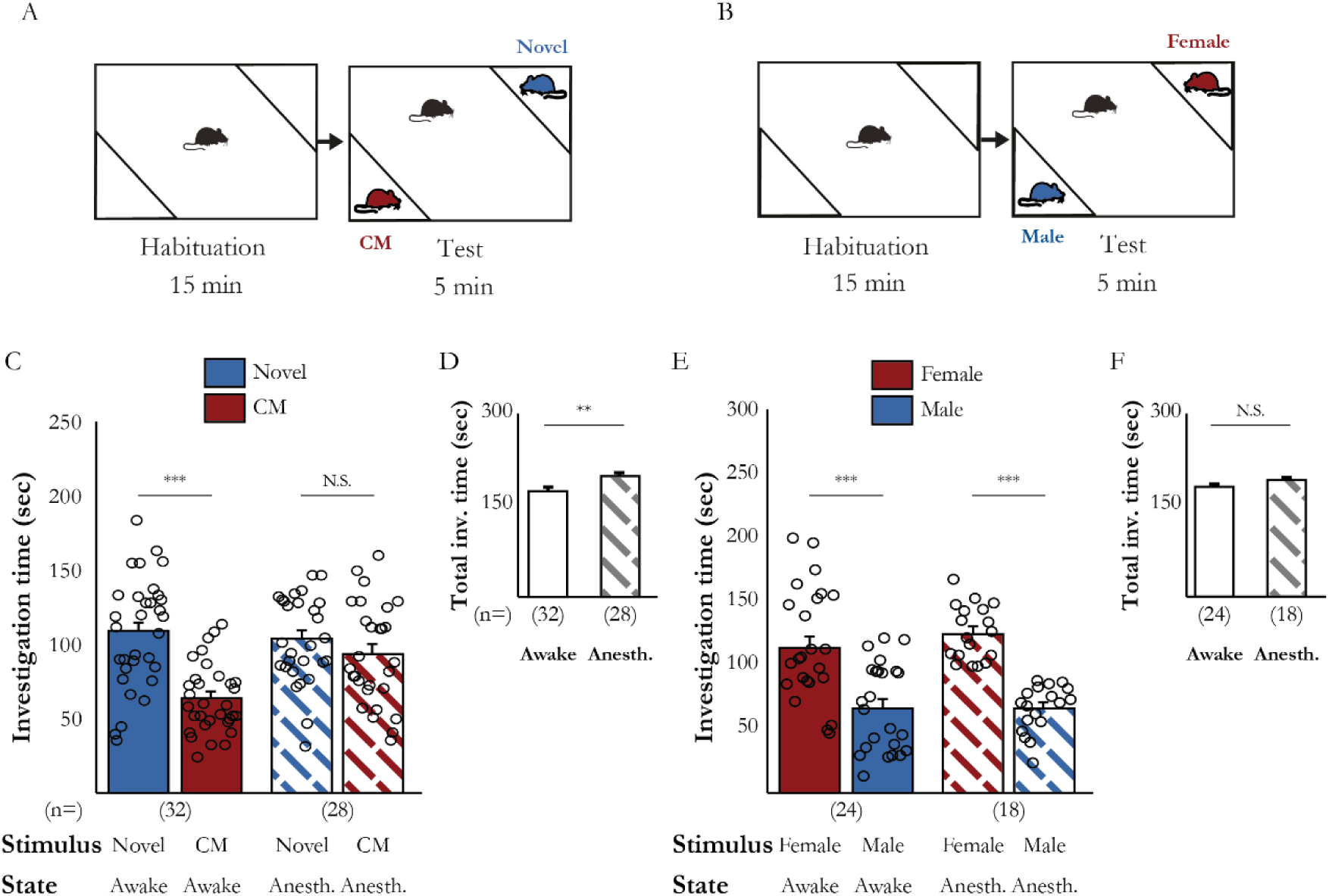
Social but not sex recognition is impaired when social stimuli are anesthetized. (A-B) Schematic descriptions of the familiarity (A) and sex (B) discrimination tests used. (C) Mean investigation time of awake (filled bars) and anesthetized (dashed bars) stimuli in the familiarity discrimination test. Number of tested subjects (n), stimulus type and state of stimulus are denoted below (Awake: t(31)=5.622, p<0.001; Anesthetized: t(27)=0.969, p=0.171, paired t-test). (D) Mean total investigation time of awake (empty bar) and anesthetized (dashed bar) stimuli in the familiarity discrimination test (Total time: t(58)=-2.479, p=0.008, independent t-test). (E-F) Similarly to C-D, for sex discrimination. (Awake: t(23)=3.596, p=0.001; Anesthetized: t(17)=-7.731, p<0.001, paired t-test; Total time: t(40)=-1.075, p=0.145, independent t-test). **p<0.01, ***p<0.001

Next, we examined whether the preference of male mice to investigate a female rather than a male mouse (termed by us sex discrimination; Fig. 1B) also depends upon the behavior of the stimuli. We found that mice preferred a female over a male even when both stimuli were anesthetized (Stimulus x Group: F(1,40)=0.410, p=0.263, mixed model ANOVA; Fig. 1E-F). Thus, at least for mice, sex discrimination does not rely on the behavior of the stimulus but rather most likely on chemosensory cues *per se*.

We then checked whether both stimuli need to be anesthetized so as to prevent discrimination between CM and novel stimuli, or whether anesthetizing only one suffices. We found that anesthetizing the CM while keeping the novel stimulus awake prevented familiarity discrimination (t(14)=0.911, p=0.189, paired t-test; Fig. 2A), whereas normal discrimination was observed in the opposite case (t(38)=3.282, p=0.001; Fig. 2B). These results suggest that anesthetizing the CM prevented its recognition by the subject, which considered the CM to be a novel stimulus. Accordingly, when two CMs, one awake and one anesthetized, were used as stimuli, the subject discriminated between them and investigated the anesthetized CM for significantly more time, as if the anesthetized CM was considered as a novel stimulus (t(18)=3.413, p=0.002, paired t-test; Fig. 2C). Such discrimination between CMs was not observed if one of them was injected with saline rather than the anesthetic (t(18)=0.844, p=0.205, paired t-test; Fig. 2D), thereby ruling out the possibility that alarm pheromones released following the injection prevented CM recognition. We, therefore, concluded that mouse and rat subjects do not recognize an anesthetized CM.

**Figure 2.**
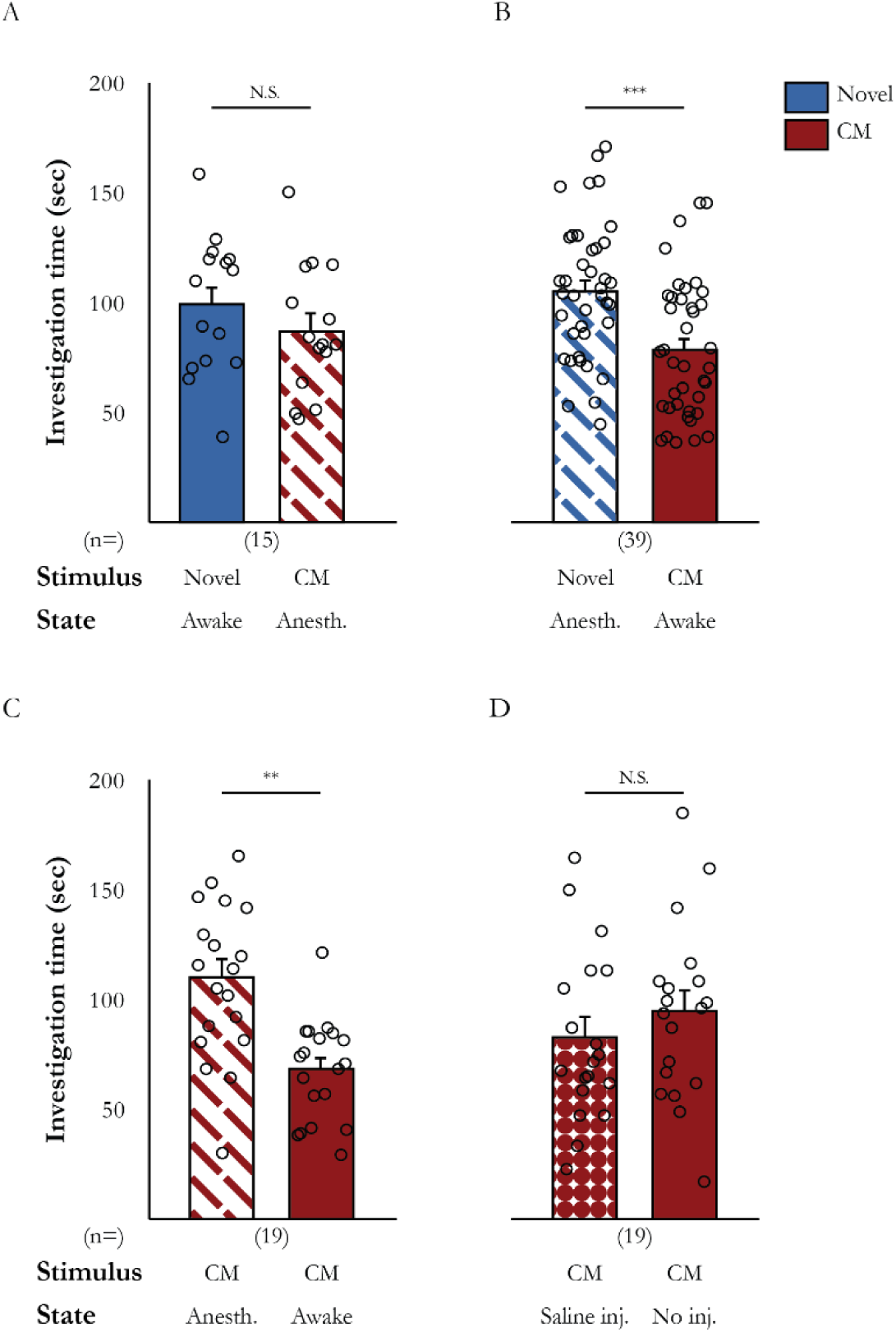
Social recognition is impaired due to a lack of activity by an anesthetized familiar stimulus. (A) Mean investigation time of both stimuli, when only the CM was anesthetized (t(14)=0.911, p=0.189, paired t-test). (B) Mean investigation time of both stimuli, when only the novel stimulus was anesthetized case (t(38)=3.282, p=0.001, paired t-test). (C) Mean investigation time of two CMs, when one was anesthetized stimulus (t(18)=3.413, p=0.002, paired t-test). (D) Mean investigation time of two CMs, when one was injected with saline (t(18)=0.844, p=0.205, paired t-test). **p<0.01, ***p<0.001

### Active behavior of social stimuli is required for subjects to learn their identity

Our results thus far can be explained by a requirement for active behavior of social stimuli for social recognition in rats and mice. An alternative explanation for our findings to this point could be that anesthesia modified the chemosensory signature of the CM, such that it could no longer be recognized by the subject. If this, however, was the case, then the subject should still be able to learn the identity of anesthetized novel stimuli and recognize them afterwards. Therefore, we used the social preference (SP)/social novelty preference (SNP) paradigm (Fig. 3A) ^34^ to determine whether mice learn to identify a social stimulus during the SP test and distinguish this stimulus from a novel stimulus during the subsequent SNP test. As is apparent from Fig. 3B, when awake stimuli were used, subject mice exhibited a clear preference towards the novel social stimulus in both the SP (upper panel, t(38)=-3.692, p<0.001, paired t-test) and SNP (lower panel, t(37)=5.137, p<0.001) tests. In contrast, when anesthetized stimuli were used for both tests (Fig. 3C), the subjects did not discriminate between the novel and familiar social stimuli in the SNP test (lower panel, t(18)=0.772, p=0.225), despite the fact that the subjects encountered the same anesthetized familiar stimulus during the SP test and investigated it normally (upper panel, t(18)=-4.870, p<0.001). This lack of discrimination between novel and familiar stimuli during the SNP test was also observed if both social stimuli were anesthetized during the SP test and awake during the SNP test (SP - t(18)=-5.811, p<0.001, paired t-test; SNP - t(18)=0.085, p=0.467; Fig. 3D). Similar results were obtained even when the SP test was replaced by free interaction with a social stimulus (Supp. Fig. 3), suggesting that a lack of free access to the anesthetized stimulus was not the reason for the lack of social discrimination in the SNP test.

**Figure 3.**
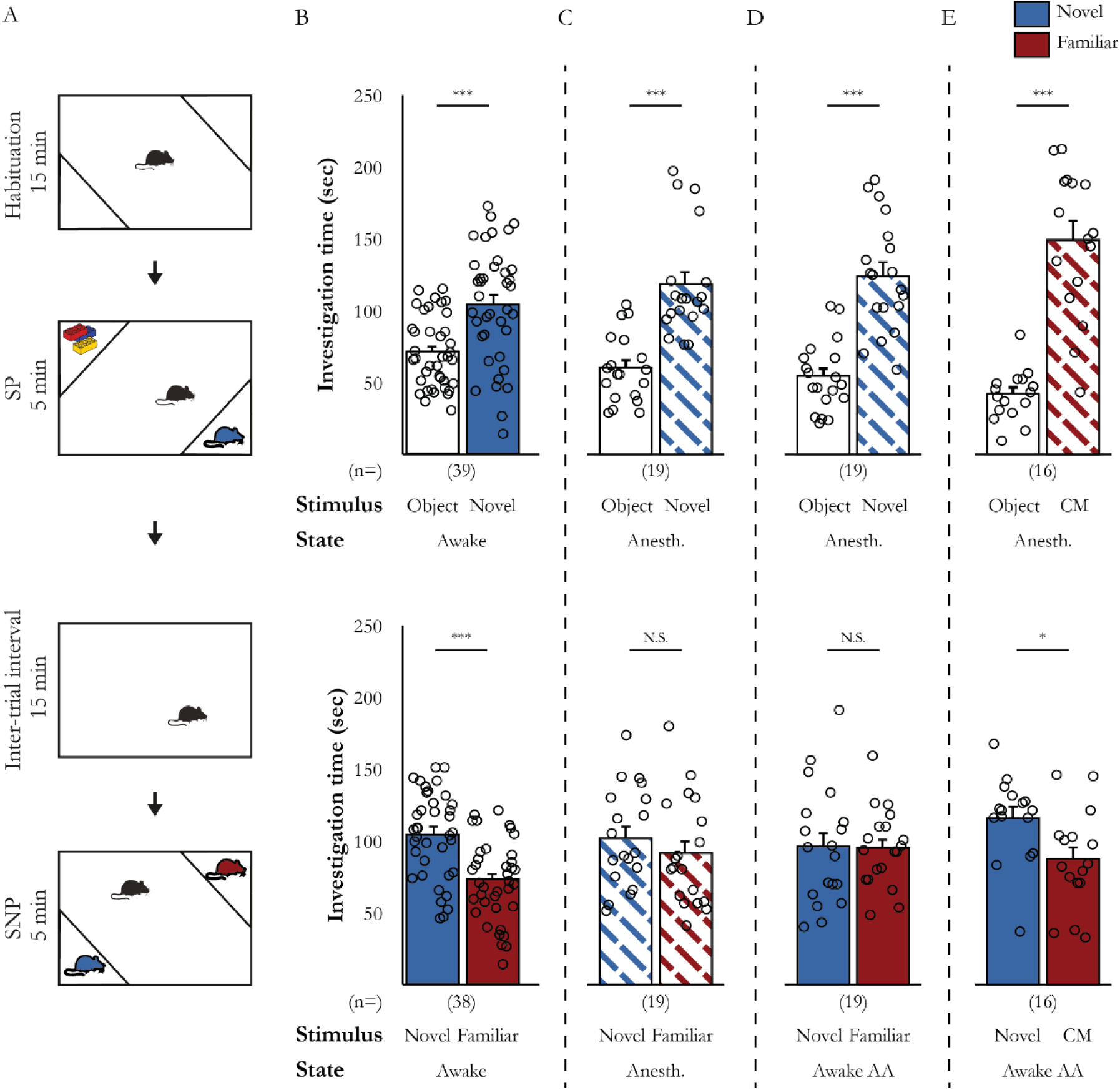
Adult male mice do not learn to recognize a social stimulus while the stimulus is anesthetized. (A) Schematic description of the SP/SNP paradigm. (B) Mean investigation times in the SP (above) and SNP (below) tests, using awake stimuli throughout (SP: t(38)=-3.692, p<0.001; SNP: t(37)=5.137, p<0.001, paired t-test). (C) As in B, using anesthetized stimuli throughout (SP: t(18)=-4.870, p<0.001; SNP: t(18)=0.772, p=0.225, paired t-test). (D) As in B, with both social stimuli were anesthetized during the SP test and were woken after anesthesia (Awake AA) during the SNP test (SP: t(18)=-5.811, p<0.001; SNP: t(18)=0.085, p=0.467, paired t-test). (E) As in D, using a CM instead of the novel social stimulus in the SP test (SP: t(15)=-6.562, p<0.001; SNP: t(15)=2.180, p=0.023, paired t-test). *p<0.05, ***p<0.001

These results suggest that not only do mice not identify an anesthetized stimulus based on their previous encounter (that took place during the SP test), they also cannot learn the identity of an anesthetized novel social stimulus. In contrast, when an anesthetized CM was used as a social stimulus during the SP test (t(15)=-6.562, p<0.001; Fig. 3E, upper panel), the subjects did discriminate between it and a novel stimulus during the following SNP test in which both stimuli were awake (t(15)=2.180, p=0.023; Fig. 3E, lower panel). This result demonstrates that if the subject is already experienced with the awake stimulus (as between CMs), the subject will recognize it even after this stimulus was anesthetized and woken. Overall, the results further support a crucial role for the behavior of the stimulus in social recognition in mice.

### Mice do not discriminate between anesthetized stimuli even following stimulus-specific social fear conditioning

Thus far, we relied on the innate social novelty preference of the animals as driving their social recognition behavior. Yet, it may be possible that while subject mice indeed discern between novel and familiar anesthetized stimuli, they do not exhibit this innate novelty-seeking tendency towards these individuals. Therefore, to confirm our conclusions, we developed a test that does not rely on the innate social novelty preference of mice, in which a behavioral paradigm of stimulus-specific social fear conditioning (SFC) was employed. In this paradigm (schematically described in Fig. 4A), we conducted two consecutive SP tests with each subject (separated by 15 min) before the fear conditioning session. For each of these tests, we used a social stimulus from a specific mouse strain (C57BL/6J and ICR; Fig. 4A upper panel) so as to enhance the ability of a subject to discriminate between these individuals. Twenty minutes after the second SP test, we conducted a 5 min SFC session using the same ICR stimulus used for the previous SP test, but in a different spatial context (Fig. 4A, middle panel). During this session, every time the subject tried to investigate the ICR mouse, it was punished by a mild (0.4 mA, 750 msec) electrical foot shock. In most cases, only 2-3 shocks were required to stop all attempts of the subject to investigate the social stimulus during this session. Twenty minutes later, we performed two SP tests with the same social stimuli as used before the conditioning session (Fig. 4A, lower panel). As apparent in Fig. 4B for awake stimuli, prior to conditioning, subject mice exhibited similar social preference to both stimuli (upper panel). However, following SFC, subjects still showed clear social preference for the C57Bl/6J stimulus, but lost their preference towards the conditioned ICR mouse (Stimulus x Time: F(3,21)=9.961, p<0.001, two-way repeated ANOVA. Fig. 4B). Thus, using this paradigm we could clearly discriminate between the two social stimuli on the basis of learned fear rather than any innate tendency.

We then considered whether subjects can discriminate between social stimuli anesthetized only during the second set of SP tests, conducted after the SFC (performed with an awake ICR stimulus). As apparent in Fig. 4C (lower panel), subjects in this case showed similar social preference towards both anesthetized social stimuli, exactly as they did before the SFC towards the same stimuli while they were awake (Stimulus x Time: F(1.839,14.713)=0.707, p=0.249, two-way repeated ANOVA. Fig. 4C). These results suggest that subject mice did not recognize the anesthetized ICR stimulus as the conditioned stimulus. Moreover, when both social stimuli were anesthetized throughout the course of the experiment (including during the conditioning session), subject mice exhibited a general SFC and still could not distinguish between the two stimuli after conditioning (Stimulus x Time: F(1.758,15.825)=17.218, p<0.001, two-way repeated ANOVA. Fig. 4D). These results further support our conclusion that discrimination between individuals of the same sex relies on their behavior and not only on chemosensory cues they passively release. Finally, similar results were obtained with SD rats using a slightly modified behavioral paradigm (Supp. Fig. 4).

**Figure 4.**
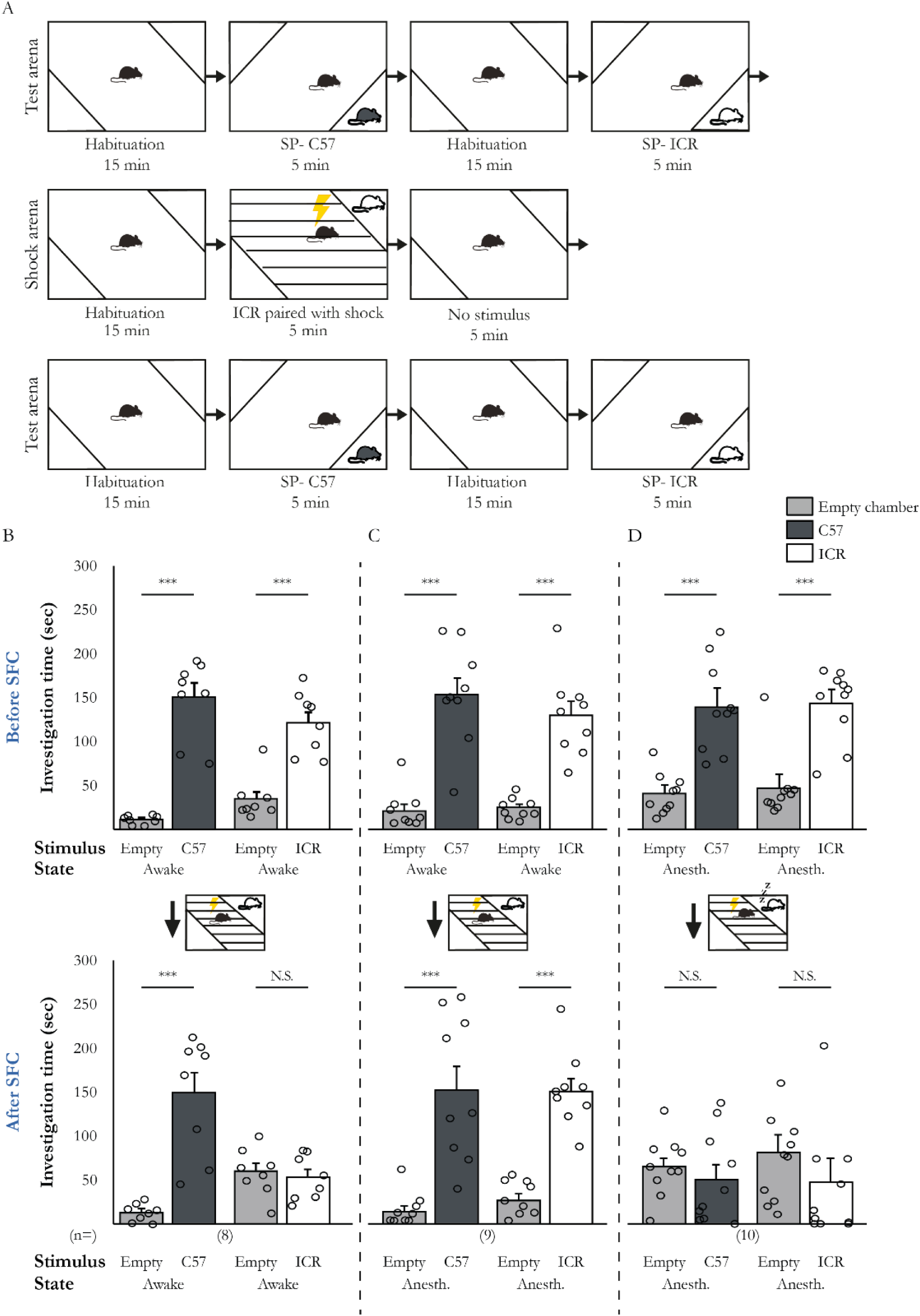
Impaired discrimination between anesthetized social stimuli following social fear conditioning (SFC). (A) Schematic description of the SFC paradigm. Two SP tests (each with a distinct stimulus) were conducted before (upper) and after (lower) the fear conditioning session (middle panel). (B) Mean investigation time of both during SP tests before (above) and after (below) SFC, using awake stimuli (before: C57BL/6J: t(7)=8.220, p<0.001, ICR: t(7)=5.104, p<0.001; after: C57BL/6J: t(7)=5.314, p<0.001; ICR: t(7)=-0.448, p=0.334, paired t-test). (C) Same as B, using stimuli that were anesthetized only after the SFC session (before: C57BL/6J: t(8)=5.372, p<0.001, ICR: t(8)=5.683, p<0.001; after: C57BL/6J: t(8)=4.463, p=0.001, ICR: t(8)=6.574, p<0.001, paired t-test). (D) Same as B, using stimuli that were anesthetized throughout the experiment (before: C57BL/6J: t(9)=-4.624, p<0.001, ICR: t(9)=-4.132, p=0.002; after: C57BL/6J: t(9)=0.63, p=0.272, ICR: t(9)=-1.009, p=0.170, paired t-test). ***p<0.001

### Social discrimination but not sex discrimination depends upon somatosensory, auditory and chemosensory cues

If behavior of stimuli is indeed required for social recognition in mice and rats, one can ask what sensory modalities are recruited for detecting it? Our behavioral experiments were performed under dim red light that is thought to be invisible to rats and mice ^35,36^ and since mice subjects did not discriminate between anesthetized C57BL/6J and ICR stimuli following SFC, despite their distinct fur color, it would seem that vision not play a central role here. However, both somatosensory and auditory sensations can be used to detect touch, movement or vocalization of the stimuli ^37^. Therefore, we examined the effects of impairing somatosensory, auditory or olfactory sensations of the subjects on both familiarity discrimination between CM and novel social stimulus, discrimination which depends upon the behavior of the stimulus (Fig. 1B), and sex discrimination, which does not (Fig. 1E). We found that impairing somatosensation by tearing the whiskers of subjects several days before the test abolished their familiarity discrimination ability (within stimulus: F(1,29)=9.562, p=0.002; Within mouse: F(1,29)=2.851, p=0.051, two-way repeated ANOVA. Fig. 5A), without affecting total investigation time (t(29)=-1.689, p=0.051, paired t-test; Supp. Fig. 5A). In contrast, whiskerless mice showed normal sex preference behavior (Stimulus x Group: F(1,42)=0.595, p=0.223, mixed model ANOVA. Fig. 5B), with a slight reduction in total investigation time (t(42)=3.853, p<0.001, independent t-test; Supp. Fig. 5B).

**Figure 5.**
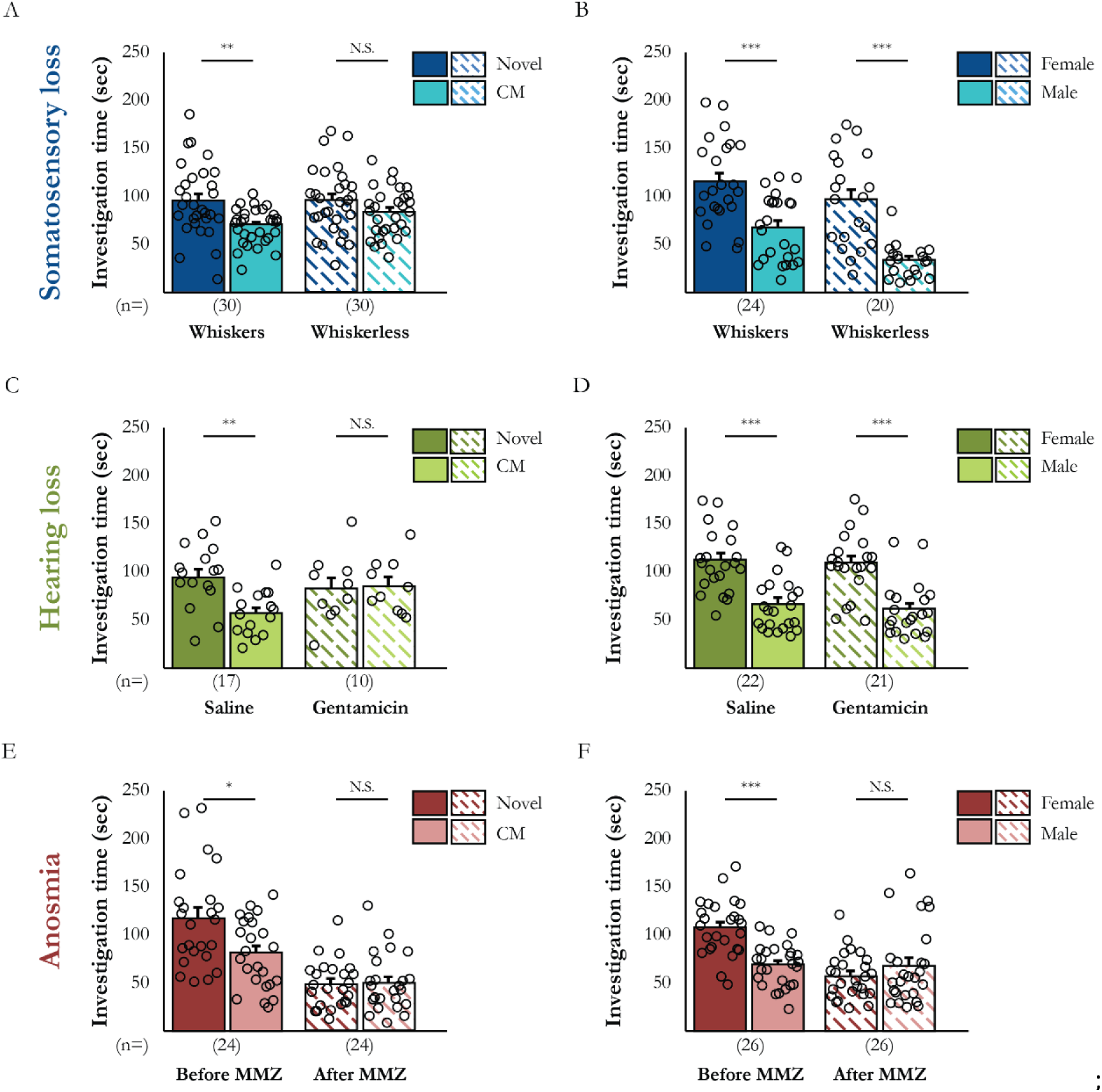
Familiarity but not sex discrimination relies on the auditory and somatosensory systems. (A) Mean investigation time of the distinct stimuli by subjects with (filled bars) and without (dashed bars) whiskers, in the familiarity discrimination test (whiskers: t(29)=3.096, p=0.002, whiskerless: t(29)=1.296, p=0.103, paired t-test). (B) As in A, for the sex discrimination test (whiskers: t(23)=3.596, p=0.001, whiskerless: t(19)=5.844, p<0.001, paired t-test). (C-D) As in A-B, for subjects with an auditory system damaged by gentamicin injection (familiarity discrimination: saline: t(16)=3.400, p=0.002, gentamicin: t(9)=-0.159, p=0.439, paired t-test; sex discrimination: saline: t(21)=4.195, p<0.001, gentamicin: t(20)=4.047, p<0.001, paired t-test). (E-F) As in C-D, for subjects with an olfactory system damaged by MMZ injection (familiarity discrimination: before MMZ: t(23)=2.323, p=0.015, after MMZ: t(23)=-0.213, p=0.417, paired t-test; sex discrimination: before MMZ: t(25)=5.742, p<0.001, after MMZ: t(25)=-1.090, p=0.143, paired t-test). *p<0.05, **p<0.01, ***p<0.001

To assess the effect of hearing loss, we examined the behavior of animals centrally injected with gentamicin, an antibiotic that kills cochlear hair cells ^38,39^ (Supp. Fig. 6), and compared this behavior to that of saline-injected animals. Like whiskerless animals, mice with hearing loss exhibited a lack of social discrimination (Stimulus x Group: F(1,25)=3.967, p=0.029, mixed model ANOVA. Fig. 5C) but not of sex discrimination behavior (Stimulus x Group: F(1,41)=0.016, p=0.451, mixed model ANOVA. Fig. 5D), with no change in total investigation time (familiarity discrimination test: t(25)=-1.307, p=0.101, independent t-test; sex discrimination: χ^2^(2)=12.151, p=0.001, Kruskal-Wallis test, *post hoc* - saline-gentamicin: U=171.5, p=0.074, saline-gentamicin+ whiskerless: U=77, p=0.012, gentamicin-gentamicin+ whiskerless: U=39, p<0.001, Mann-Whitney U test; Supp. Fig. 5D-E).

To examine the effects of anosmia, we compared the behavior of animals before and after a single injection of methimazole (MMZ), a drug used for treating hyperthyroidism and which is known to kill olfactory sensory neurons ^40,41^ (Supp. Fig. 7). As apparent in Fig. 5E-F, MMZ-induced anosmia abolished not only the social discrimination ability of the animals (Stimulus x Group: F(1,46)=4.858, p=0.017, mixed model ANOVA. Fig. 5E) but also their sex preference behavior (Stimulus x Group: F(1,25)=15.981, p<0.001, two-way repeated ANOVA. Fig. 5F). Moreover, anosmia also caused a significant reduction in total investigation time in both tests (familiarity discrimination: t(46)=7.980, p<0.001,independent t-test; sex discrimination: t(25)=4.629, p<0.001, paired t-test; Supp. Fig. 5G-H), suggesting a reduction in general motivation for social interaction. Thus, familiarity discrimination seems to require not only intact olfaction but also functional hearing and whisker-dependent somatosensation. In contrast, sex discrimination relies on olfaction but not on hearing or whisker-dependent somatosensation, in accordance with our previous results using anesthetized stimulus-presenting mice (Fig. 1).

### Somatosensory and auditory cues can be used redundantly for social recognition

We further examined how the aforementioned sensory impairments affect the ability of sensory-deprived subjects to learn the identity of a novel social stimulus-presenting individual in the SP/SNP paradigm presented above (Fig. 3). Surprisingly, we found that both whiskerless and gentamicin-injected subjects showed significant preference for investigating the novel social stimulus in the SNP test (somatosensory: Stimulus x Time: F(1,27)=0.023, p=0.440, two-way repeated ANOVA; auditory: Stimulus x Group: F(1,41)=2.343, p=0.067, mixed model ANOVA. Fig. 6A-B), while MMZ-injected animals did not show such a preference (Stimulus x Time: F(1,15)=4.652, p=0.024, two-way repeated ANOVA. Fig. 6C). However, when a new cohort of subjects were subjected to both somatosensory and hearing impairment (i.e., whiskers tearing and gentamicin injections), they could not discriminate between the two stimuli in the SNP test (Stimulus x Group: F(1,33)=3.313, p=0.039, mixed model ANOVA. Fig. 6D), while preserving intact sex-reference behavior (Supp. Fig. 8). Thus, it seems as if somatosensory and auditory cues are required for social recognition in a redundant manner, whereas olfactory cues are a pre-requisite.

**Figure 6.**
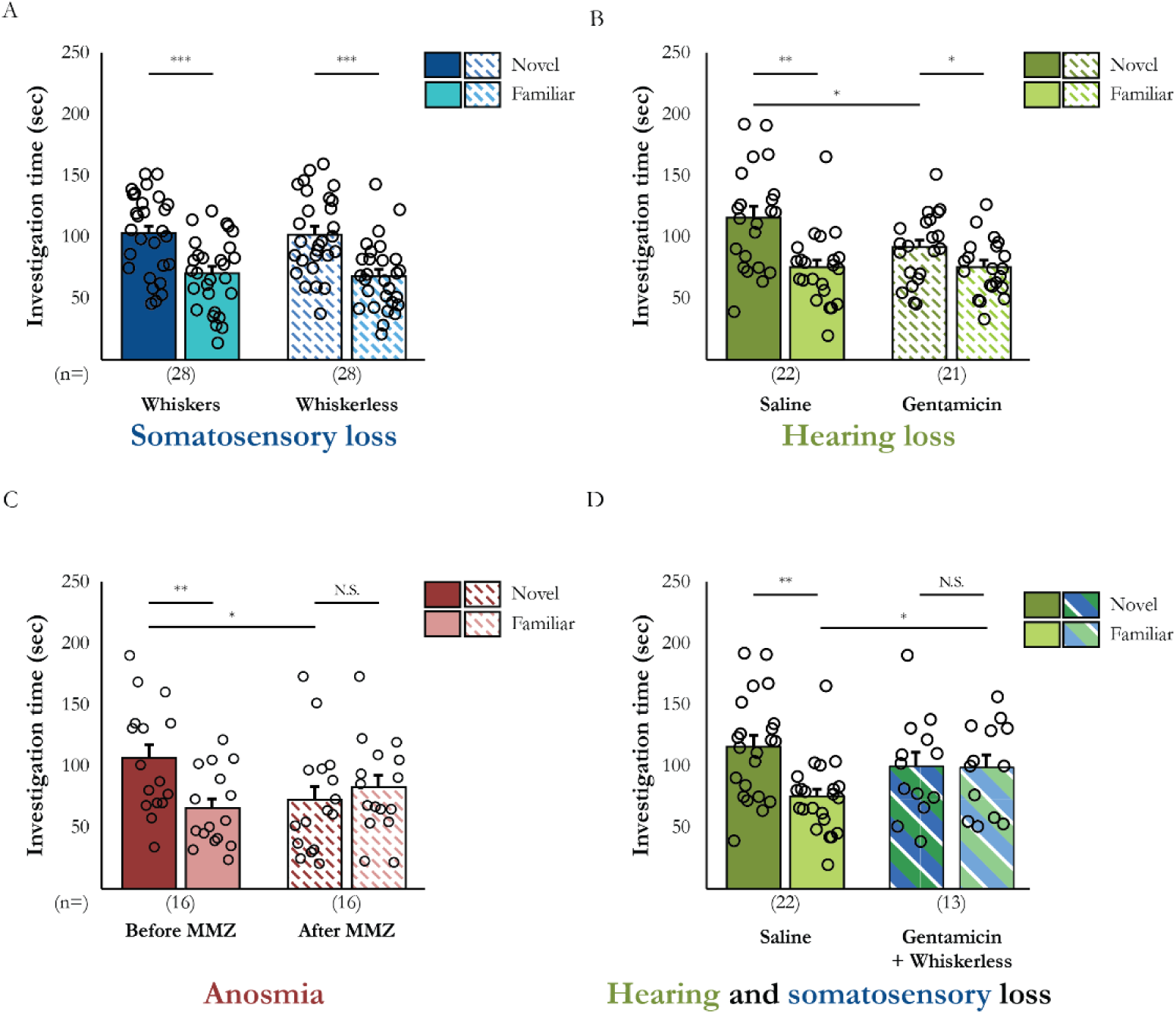
Either the auditory or somatosensory systems can be used for learning the identity of a novel social stimulus. (A) Mean investigation times of both stimuli in the SNP test, as performed by control and whiskerless subjects (whiskers: t(27)=4.239, p<0.001, whiskerless: t(27)=4.017, p<0.001, paired t-test). (B) As in A, for subjects injected with saline or gentamicin (Saline: t(21)=2.959, p=0.004; Gentamicin: t(21)=2.054, p=0.027, paired t-test). (C) As in A, for subjects injected with saline or MMZ (before MMZ: t(15)=2.637, p=0.010, after MMZ: t(15)=-0.609, p=0.276, paired t-test). (D) As in A, for subjects with damage in both the somatosensory and auditory systems (saline: t(21)=2.959, p=0.004, gentamicin+ whiskerless: t(12)=0.038, p=0.486, paired t-test). *p<0.05, **p<0.01, ***p<0.001

### Movement of stimuli can produce somatosensory and auditory cues required for social recognition

The results presented so far suggest that both the auditory and somatosensory modalities are involved in social discrimination in a redundant manner. We, therefore, looked for stimulus-generated cues that may be detected by both modalities. A primary candidate is the movement of the stimulus, as movement produces both somatosensory (via the whiskers) and auditory cues, and hence may be detected by both modalities and supply information required for social recognition. To examine if there are any differences in the movements generated by a CM and a novel social stimulus during the familiarity discrimination test, we used a novel movement monitoring system comprising an array of piezo-electric sensors placed at the floor of the triangular chambers containing the social stimuli (Fig. 7A-B). We then recorded the electrical signals generated by the sensors, which reflect the movement of each stimulus, during familiarity discrimination tests performed by 35 male C57BL/6J subjects. The raw piezo signal trace recorded along the time course of each session (see example in Fig. 7C) was normalized to the peak signal separately for each chamber so as to correct for differences in mass and strength among the various stimuli. We subsequently quantified the number of major movements, defined by peaks that crossed a threshold ranging between 10-30% of the maximal signal. We found that at all threshold levels, novel stimuli generated significantly higher numbers of major movements, especially during the first 2-3 minutes of the test (threshold: 10%: χ^2^(9)=58.353, p<0.001, Friedman test; *post hoc* - min1: Z=-2.842, p=0.002, min2: Z=-2.424, p=0.008, min3: Z=-2.703, p=0.003, min4: Z=-1.458, p=0.072, min5: Z=-1.097, p=0.136, Wilcoxon signed rank test; 20%: χ^2^(9)=44.257, p<0.001, Friedman test, *post hoc* - min1: Z=-2.138, p=0.016, min2: Z=-1.941, p=0.026, min3: Z=-1.778, p=0.037, min4: Z=-0.475, p=0.317, min5: Z=-0.205, p=0.418, Wilcoxon signed rank test; 30%: χ^2^(9)=27.26, p<0.001, Friedman test, *post hoc* - min1: Z=- 1.760, p=0.039, min2: Z=-1.582, p=0.056, min3: Z=-0.915, p=0.180, min4: Z=-0.299, p=0.382, min5: Z=-0.655, p=0.256, Wilcoxon signed rank test; Fig. 7D-F). Thus, during the early phase of the social discrimination test, novel stimuli seemed to be more active than were CMs. To examine if the behavior of the subject was affected by the movement of the stimuli, we measured the time spent by the subject investigating each of the stimuli following a major movement made by that individual (using a threshold level of 25%), as compared to when no major movement occurred. We found that at resting conditions (i.e., after a 4-s break in the investigation behavior), subject mice showed a reduction in their likelihood to restart investigating the CM following it making a major movement (Fig. 7G, upper panel, red line), as compared to periods when no major movement was observed (green lines). No such tendency was found towards novel social stimuli, with differences in periods of movement or no movement were found to be statistically significant (CM: Z=-3.513, p<0.001, Novel: Z=-0.915, p=0.180, Wilcoxon signed rank test. Fig. 7H). As a control, we made the same calculations for periods that followed a 4 sec window of investigating a stimulus and found no effect of major movements in this case (CM: Z=-0.231, p=0.485, Novel: Z=-0.556, p=0.289, Wilcoxon signed rank test. Fig. 7G-H, lower panels). To confirm that these differences were not caused by a preference of the subjects for investigating a novel stimulus, we analyzed the results separately for subjects who preferred the novel stimulus (Fig. 7I, upper panel) and those who preferred the CM (Fig. 7I, lower panel). We found no differences between the groups (Supp. Fig. 9). Thus, it seems that there is a specific negative effect of CM movement, but not the movement of a novel stimulus, on subject motivation to start investigating. Overall, these results reveal that familiar and novel stimuli exhibit different movement patterns in the familiarity discrimination test, and that subjects react to these patterns in a stimulus-dependent manner. These results may thus explain why subject mice failed to recognize CMs when they were anesthetized and could not move.

**Figure 7.**
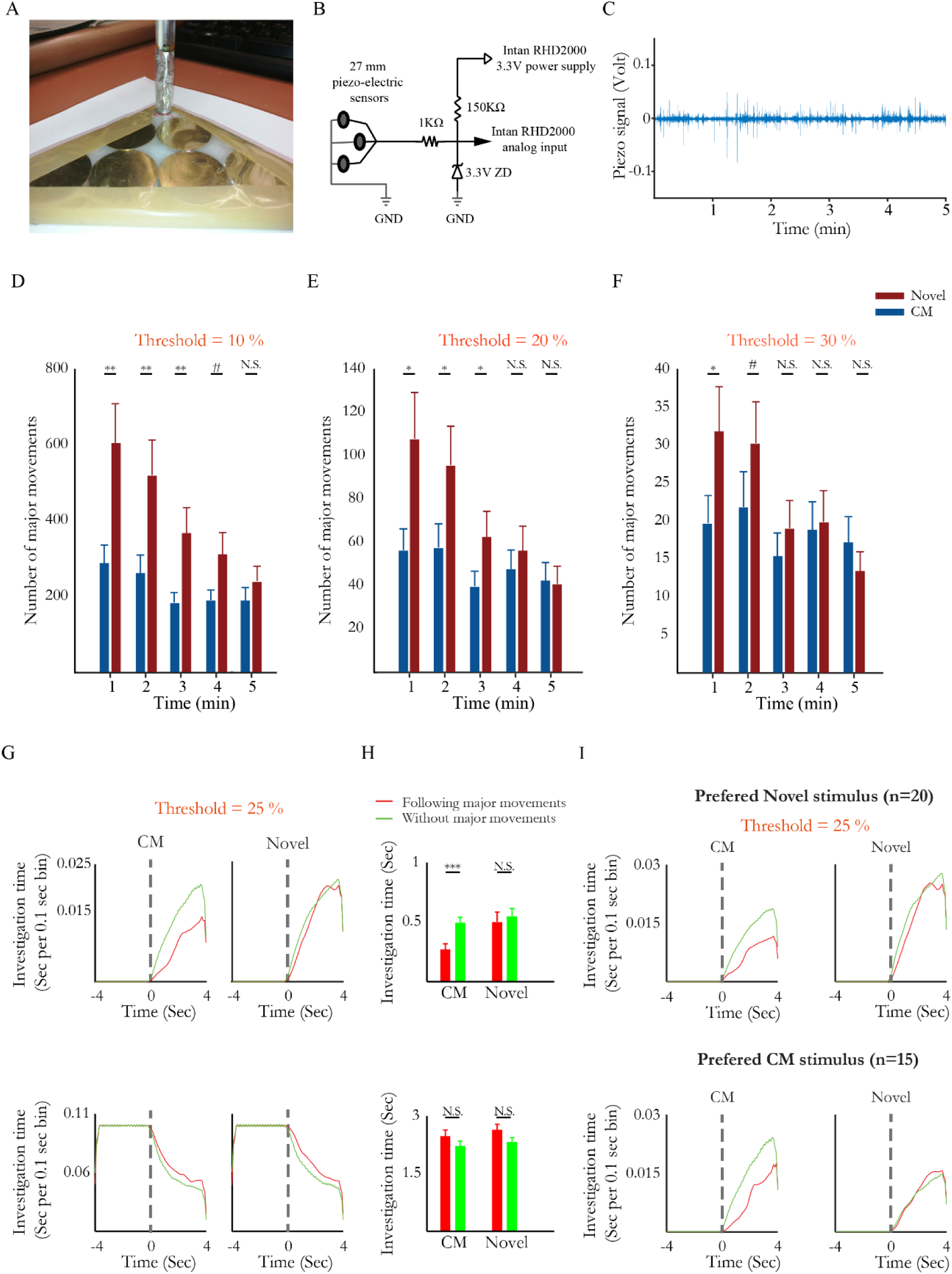
Differential movement by social stimuli during familiarity discrimination draws distinct subject responses. (A) Picture of the piezoelectric sensors array lining the floor of the stimulus chamber. (B) Schematic description of the electric circuit used for connecting the sensors. (C) Raw electrical signal recorded during an experiment from one of the stimuli. (D) Mean number of major movements made by each of the stimuli, using a threshold of 10% of the maximal signal. (E) Same as D, using a threshold of 20%. (F) Same as D, using a threshold of 30%. (G) Above - mean investigation time of the distinct stimuli, after 4-sec period with no investigation by the subject, with (red) and without (green) major movement made by the CM (left) or novel social stimulus (right). Time 0 marks the beginning of movement. Below – same as above, following 4-sec period of stimulus investigation. (H) Statistical analysis of the results shown in G. (I) Above – mean investigation times of the stimuli, after a 4 sec period of no investigation, with (red) and without (green) major movement made of the stimulus, for subjects that preferred the novel social stimulus over the CM. Below – as above, for animals that preferred the CM. #p<0.1, *p<0.05, **p<0.01

Altogether, the results of this study suggest that subject mice integrate chemosensory and movement cues, acquired via multiple sensory modalities, including the olfactory, auditory and somatosensory modalities, to discriminate between same-sex stimuli.

## Discussion

In this study, we explored the role of stimulus behavior in social recognition in laboratory rats and mice. We hypothesized that social recognition in these animals does not depend solely on chemosensory signature passively emitted by the stimulus (i.e., the signaler) and detected by the subject (i.e., the receiver), in analogy to face recognition by humans. Instead, social recognition involves detection of behavioral cues, thus requiring both activity of the stimulus and integration of information arriving via several sensory modalities by the subject. To test this hypothesis, we examined two types of social recognition, discrimination between conspecifics based on their sex (i.e., sex discrimination) and discrimination between conspecifics based on previous encounters (i.e., familiarity discrimination). Using the unique design of our experimental system, we compared sex and familiarity discrimination by subject animals when challenged with sets of awake and anesthetized stimuli. We found that adult male mice could discriminate between male and female stimuli even when these individuals were anesthetized, suggesting that chemosensory signals passively emitted by the stimuli were sufficient for sex discrimination. This conclusion, which is in accordance with multiple previous studies ^42,43^, is further supported by the observation that olfactory impairment was the only treatment that affected sex discrimination, whereas somatosensory and auditory impairment had no influence on this behavior. In contrast, both mice and rats failed to discriminate between familiar and novel social stimuli, either if these animals were anesthetized or following impairment of each of the sensory modalities in the subjects considered, suggesting a role for the behavior of the social stimulus, as acquired via multiple sensory modalities, in familiarity recognition. These findings thus confirm our hypothesis.

### Working with anesthetized stimuli

The number of behavioral studies exploring social recognition that used anesthetized stimuli is surprisingly small. Anesthetized same-sex conspecifics were found to elicit defensive responses and ultrasonic vocalizations in rats ^44,45^. Latané and Glass ^46^ reported a reduction in the level of contacts made by rats with anesthetized stimuli as compared to freely moving stimuli and concluded that movement is important for the attractiveness of social stimuli. It should be noted that in all of the experiments we conducted, no changes were noted in the general time that subject rats and mice dedicated to investigating their anesthetized counterparts, as compared to awake animals, suggesting that no reduction in attraction to anesthetized stimuli had occurred. Johnston and Peng ^47^ used the cross-odor paradigm with male hamsters to show that physical contact with a female body is required for cross-odor habituation to the scent of that female. These authors further concluded that hamsters rely on multimodal integration for social recognition, similarly to what we now report for rats and mice. However, this earlier study found that contact with anesthetized female was as effective as was contact with awake female for this type of recognition, suggesting no role for the behavior of the stimulus in this type of intersexual social recognition. Thus, hamsters may differ from rats and mice in terms of the type of multimodal cues used for social recognition. Notably, a recent study using a novel methodology for assessing social investigation over a 100 min period found that adult male mice investigated anesthetized CMs more than they did novel conspecifics, with each being encountered separately. However, this difference was observed only after 10 min of exposure to the stimulus; no difference was found during the first 10 min, in accordance with our results. Thus, it may be possible that after prolonged periods of exposure to anesthetized stimuli mice can distinguish between such individuals. Seemingly, this does not occur during the widely used social discrimination tests lasting less than 10 min.

One concern regarding our use of anesthetized animals is the possibility that the anesthesia procedure we employed changed the odor of the social stimulus, thus interfering with subject recognition of animals previously encountered prior to the anesthesia. Multiple observations from our experiments, however, argue against this possibility. First, subjects did not differentiate saline-injected from non-injected CMs, suggesting no interference by injection-induced release of alarm pheromones. Second, the anesthesia had no effect on the ability of the subjects to discriminate between male and female stimuli, suggesting that such treatment did not mask chemosensory cues emitted by these individuals. Third, subjects did not learn to recognize an anesthetized stimulus even if this individual was still anesthetized during the discrimination test, thus ruling out the possibility that anesthesia induced a novel chemosensory signature the social stimulus. Finally, the full agreement of the results of sensory impairment experiments, all conducted with awake stimuli, with the results of social discrimination experiments conducted with anesthetized stimuli, suggests that all the results reflect the same dependence of familiarity discrimination on cues generated by the behavior of social stimuli.

### Social recognition by chemosensory cues

A large body of evidence suggests that chemosensory cues can alone mediate several types of social recognition ^48,49^, such as mate and kin recognition ^25,50-54^. In the case of familiarity recognition, the picture is less clear. Several studies have shown that anosmic rats and mice lose their ability to discriminate between familiar and novel conspecifics ^30,55-57^. These results, which are in agreement with the data presented here, does not rule out the involvement of other sensory modalities in familiarity discrimination. Multiple other studies have used operant conditioning to demonstrate that rats and mice can learn to discriminate between odors of social stimuli even if the animals involved are almost genetically identical ^6,58,59^. Operant conditioning is, however, well known for revealing astonishing capabilities of discrimination between almost identical complex sets of signals. For example, mice can be trained to discriminate between stimuli on the basis of activity of a single cortical neuron ^60^. Such skills may define the limits of learning capabilities, yet are not necessarily employed in natural conditions. Other studies involving more ethologically relevant habituation-based learning showed that small rodents can discriminate between chemosensory cues derived from individual conspesifics based on either the highly polymorphic major histocompatibility complex (MHC) or a combinatorial repertoire of major urine proteins (MUPs) ^17,61,62^. However, these studies largely focused on body odors, such as urine, instead of the social stimulus itself. As the mechanisms employed for recognition may differ in the context of encountering an individual conspecific than when encountering odors derived from the same individual, it is hard to draw conclusions regarding the sensory modalities involved in recognition of the actual conspecifics from these experiments ^3^.

### Using social fear conditioning for social discrimination

In the present study, we employed two distinct habituation-based tests to examine the role of behavioral cues in familiarity recognition. In the social discrimination test, we examined discrimination between a novel stimulus and a CM, while in the SP/SNP paradigm we examined discrimination between a novel stimulus and a stimulus encountered 15 minutes earlier in the same spatial context. In both cases, subjects did not associate between the same social stimulus in the awake and anesthetized states, and did not habituate to an anesthetized stimulus. These results suggest that a familiar individual that is not behaving in an active manner is not recognized, regardless of the context of the previous encounter, whether in the home cage during daily life or in the same temporal and spatial context in which the discrimination test was conducted. Habituation-based tests, however, rely on the innate tendency of rodents to explore novel stimuli more than familiar stimuli. To rule out the possibility that by anesthetizing the social stimuli we interfered with this innate tendency rather than with recognition of the stimulus, we used the novel paradigm of SFC.

The ability of SFC to overcome the innate attractiveness of social interactions in small rodents was recently demonstrated by several laboratories ^63,64^. Although these studies used SFC to induce general avoidance of social stimuli, we, nevertheless, developed a protocol that enables induction of avoidance behavior towards a specific individual, allowing other conspecifics to retain their attractiveness. We used this protocol to show that rats and mice did not discriminate between fear-conditioned and neutral social stimuli if these animals were anesthetized either during the conditioning session or during the discrimination test, despite the unpleasant experience associated with the conditioned stimulus. Notably, a previous study employed a similar approach involving aversive conditioning to female odors showed that *TrpC2* knockout mice could be conditioned to avoid female stimuli, despite their lack of innate discrimination between male and female stimuli ^65^. Therefore, our SFC results clearly support the conclusions that mice and rats do not discriminate between anesthetized stimuli due to recognition failure, rather than because of problems with motivation.

### Sensory modalities that mediate familiarity recognition

To examine which sensory modalities are involved in detection of behaviorally induced social cues that contribute to familiarity recognition, we used established methods to induce impairments in either the olfactory, somatosensory or auditory modalities of subject mice. We found that impairing any of these modalities impaired social discrimination between a novel social stimulus and a CM. The fact that only olfactory impairment interfered with sex recognition suggests that our treatments were specific to the targeted modalities and did not induce a general behavioral state leading to a lack of discrimination between stimuli. Nevertheless, when animals affected by the same impairments were examined using the SP/SNP paradigm, it was found that compromising either the auditory or somatosensory modality alone did not abolish familiarity recognition, as did olfactory impairment. Instead, we needed to impair both modalities in the same animal to prevent familiarity recognition. This can be explained by the fact that the subject was habituated to its CMs before treatment, and hence could not recognize them without these sensory modalities. In contrast, during the SP/SNP paradigm, the subject was habituated to the stimulus after treatment without the proper function of one modality (either the somatosensory or auditory modality) and thus could detect the same behavioral cues during the test using the other intact modality. Nevertheless, these results suggest redundancy between somatosensory and auditory cues.

Multiple seminal studies from the Brecht laboratory showed that social interactions of rats involve intensive bouts of facial touch ^66^, which are temporally coordinated with ultrasonic vocalizations emitted by the subjects ^37^. They also demonstrated that such facial contacts trigger social-specific responses of single units in the somatosensory cortex ^67,68^, while also modulating neuronal responses to social vocalizations in the auditory cortex ^37^. These studies suggest the existence of a substrate for the integration of sensory cues emitted by rats during close social interaction by the somatosensory, auditory and other cortices ^69^. However, unlike rats, C57BL/6J mice do not normally emit ultrasonic vocalizations during male-male interactions ^70^, an observation replicated by us in the experiments described above. Thus, we hypothesized that in mice, movements by social stimuli can supply cues that are detected by both the somatosensory and auditory modalities, both during close contact and remote interactions between the animals. We, therefore, applied a novel experimental system based on piezoelectric sensors, which produces electrical signals that are proportional to the movement of the social stimulus within its chamber. Using this system, we found that novel stimuli are more active in the chamber during the discrimination test than are familiar stimuli. We also found that subject mice responded to major movements of the stimulus in a stimulus-dependent manner. While these results do not prove that the differential movement of the stimuli is the basis for their recognition by the subject, they do show that such behavior can serve as a source of behavior-dependent social cues, which contribute to familiarity recognition in mice.

### Summary

Overall, our results contradict the widely accepted view of social recognition in rats and mice as being solely based on chemosensory cues. Instead, we suggest that some aspects of social recognition, such as familiarity recognition, require integration of information from several sensory modalities, including the somatosensory and auditory modalities. As such, social recognition in laboratory rats and mice may differ from what occurs in humans, where social recognition can be mediated by a single sensory modality, such as visual facial cues. This difference may be relevant not only to the brain systems involved in social recognition but also to the type of interactions required. While humans can easily recognize passive social stimuli, mice and rats may need to actively engage such individual so as to determine their identity. These differences may dictate fundamentally distinct types of social interactions and relationships between humans and small rodents. As laboratory mice and rats are widely used as models of human social behavior, especially in the field of neurodevelopmental disorders, such differences call for consideration.

## Acknowledgments

This study was supported by The Human Frontier Science Program (HFSP grant RGP0019/2015), the Israel Science Foundation (ISF grants #1350/12, 1361/17), by the Milgrom Foundation and by the Ministry of Science, Technology and Space of Israel (Grant #3-12068).

## Materials and Methods

### Animals

Both mice and rats were commercially obtained (Envigo, Israel). Mice subjects were naïve C57BL/6J adult (8-15 week-old) male mice, while stimuli mice were C57BL/6J juvenile (21-30 day-old) male mice, naïve adult male and female C57BL/6J mice and ICR (CD1) male mice. Mice were housed in groups of 2-5 per cage on a 12 h light/12 h dark cycle, with lights being turn on at 7 p.m. each night. Rats subjects were adult (10-15 week-old) Sprague Dawley (SD) males, while stimuli were juvenile (21-30 day-old) SD male rats, adult SD male and female rats and adult male Long Evans (LE) rats. Rats were kept in groups of 2-5 animals per cage, on a 12 h light/12 h dark cycle, with lights being turned on at 9 p.m. each night. All animals had *ad libitum* access to food (standard chow diet; Envigo RMS, Israel) and water. Behavioral experiments were performed during the dark phase, under dim red light. All experiments were performed according to the National Institutes of Health guide for the care and use of laboratory animals, and approved by the Institutional Animal Care and Use Committee of the University of Haifa.

### Anesthesia

Mice were anesthetized with an intraperitoneal (i.p.) injection of a mixture of 100 mg/kg ketamine (100 mg/ml Clorketam, v’etoquinol) and 0.8 mg/kg medetomidine (1 mg/1 ml Domitor, Orion Pharma) in sterile saline (0.1 ml mix anesthesia/10 g BW). At the end of the experiments, mice were awakened with an injection of 0.1 ml/10 g BW atipamezole (4 mg/kg) in sterile saline (5 mg/ml Antisedan). Rats were first lightly anesthetized in a ventilated box with a few drops of 99.9% isoflurane, followed by subcutaneous injections of both 0.5 mg/kg medetomidine (0.05 ml/100 g BW) and 100 mg/kg ketamine (0.1 ml/100 g BW).

### Experimental setups

#### Social discrimination

The experimental setups used for all types of social discrimination in both mice and rats (Supp. Fig. 1) have been previously described in detail ^71^. Briefly, each setup consisted of a plexiglas arena placed in the middle of an acoustic chamber. Two plexiglas triangular chambers were placed in two randomly selected opposing corners of the arena, in which an animal or an object (plastic toy) stimulus was placed. A metal mesh placed at the bottom of the triangular chamber allowed direct interaction with the stimulus through the mesh. A high-quality monochromatic camera (Flea3 USB3, Point Grey) equipped with a wide-angle lens was placed at the top of the acoustic chamber and connected to a computer, enabling a clear view and recording of subject behavior using commercial software (FlyCapture2, Flir).

#### Social fear conditioning

##### Mice

The setup used for social fear conditioning in mice was a custom-made white plexiglas arena similar in size to the main experimental setup (37 × 22 × 35 cm) but with a metal grid floor (H10-11M, Coulbourn Instruments) connected to an electrical shock-delivering unit (precision regulated animal shocker H13-14, Coulbourn Instruments). The unit was modified to deliver a single pulse of 750 ms when manually triggered.

##### Rats

The setup used for social fear conditioning in rats was a custom-made black matte plexiglas arena (78 × 27 × 51 cm) with a metal grid floor (54 × 27 cm) in the middle (H10-11R, Coulbourn Instruments). Two black rectangular plexiglas chambers (12 × 27 × 51 cm) were placed on either side of the metal grid. Interactions were achieved through a metal mesh (25 × 7 cm, 2.5 × 1 cm holes) glued to the bottom of the chambers. The grid was connected to the electrical shock-delivering unit, modified to deliver a single 750 ms pulse when manually triggered.

### Behavioral paradigms

#### Familiarity discrimination paradigm

The social discrimination paradigm consisted of 15 min habituation of subject mice to the arena containing two empty chambers. Simultaneously, stimuli mice were introduced into chambers outside the arena for acclimation. After habituation, the social discrimination test was conducted for 5 min when the subject was simultaneously introduced to social stimuli (one CM and one novel stimuli) placed at opposite corners of the arena. When anesthetized stimuli were used, such individuals were anesthetized 15 min before the test and were kept anesthetized until the end of the experiment. The stimuli received an additional dose of the anesthetic (33% of the initial dose) when signs of awakening appeared (i.e., whisker movement). Between experiments, stimuli were kept on a heating pad at 37°C. At the end of an experiment, the stimuli were awakened by an injection of atipamezole and placed on the heating pad until fully awake.

#### Sex preference test

The sex preference test consisted of 15 min habituation of subject mice to the arena containing two empty chambers. Simultaneously, stimuli mice (adult male and female) were introduced into two additional chambers outside the arena for acclimation. After habituation, the social discrimination test was performed for 5 min when the subject was simultaneously introduced to the social stimuli found at opposite corners of the arena.

#### Social Preference/Social Novelty Preference paradigm

The SP/SNP paradigm was previously described in detail ^34,71^. Briefly, the paradigm involved a 15 min window of subject mice habituation to the arena presenting two empty chambers. Thereafter, social and object stimuli were randomly introduced to distinct corners of the arena, with the SP test being performed for 5 min. Upon termination of the SP test, the chambers housing the stimuli were removed from the arena, and the subject was left alone for 15 min. Then, the chambers were returned, this time to the other two corners of the arena, with one containing the same social stimulus used in the SP test (now as a familiar stimulus) and the other containing a novel one. At this point, the SNP test was performed for 5 min. Notably, the familiar stimulus was always placed in a different corner, relative to its position in the SP test. At the end of the SNP test, the subject was returned to its home cage, while stimuli were either left in the chambers for additional experiments or returned to their home cages.

#### Social Fear Conditioning paradigm with mice

The SFC paradigm with mice consisted of 15 min habituation of the subjects to the arena presenting two empty chambers, followed by two consecutive SP tests with two distinct strains (C57BL/6J and ICR), separated by a 15 min interval. Thereafter, the subject was transferred to the social fear conditioning arena for 15 min habituation, followed by 5 min of the SFC procedure, in which the subject received a mild electrical foot shock (0.4 mA, 750 mSec) each time it tried to interact with the stimulus chamber (ICR strain). Five min after conditioning, the subject was returned to the experiment arena for 15 min habituation and two more consecutive SP tests, performed as before conditioning.

#### Social Fear Conditioning paradigm with rats

For the SFC paradigm with rats, a strain preference test was first performed on day one. The test was initiated with a 10 min habituation of the subject rats to the experimental arena presenting two empty chambers. The preference test was next conducted by introducing two novel stimuli (adults SD and LE males) for 5 min. The SFC in rats was conducted over the course of two additional days. On the first day, subject rats were allowed to investigate the conditioning arena presenting two empty rectangular chambers for 10 min. Thereafter, the two stimuli rats (LE and SD) were introduced into the black rectangle chambers found on the sides. The rats were allowed to investigate the stimuli for 5 min, and were given a 750 ms electrical foot shock (0.7mA) each time they investigated the LE stimulus. Following a 1 h interval in their home cage, the subject rats were returned to the conditioning arena for another SFC session, this time with the stimuli on the opposite sides. A day later, the subject rats were placed in the experimental arena for another strain preference test with LE and SD stimuli rats.

### Behavioral data analysis

Data analysis was performed using our custom-made TrackRodent software, as previously described ^34,71^.

### Measuring movement of social stimulus using piezoelectric sensors

#### Setup

Movements of stimuli animals were measured using six piezoelectric ceramic discs (27 mm in diameter) connected in parallel. The discs were evenly distributed along the triangulated plexiglass floor and adjusted using lamination foil. Signal from the piezo-discs were transferred to the analog input of a RHD2000 recording system (Intan Technologies) through a protective metal tube fixed to the inner wall of the triangulated chamber (see scheme in Fig. 7A, B).

#### Analysis

All signals were analyzed using a custom-made MATLAB analysis program. Raw signals were recorded at 20 kHz. Signals were then down-sampled to 2000 Hz and band-pass filtered between 10-100Hz using a Butterworth filter. Large movements were detected using a threshold of more than 20% of the maximum signal absolute value. Varying this threshold between 10 and 40% did not change the final results. For detecting a subject’s tendency to investigate a stimulus animal after movement of the latter, we analyzed all periods meeting the criteria of no social investigation by the subject and no large movements by the stimulus for at least 4 sec before a given movement by that animal (Fig.7G, H). Varying this period between 2-8 sec did not change the final results. For statistical analysis (Fig. 7H), the total investigation time within 4 sec window after the movement was considered for calculating the mean investigation time.

#### Modality impairment

##### Whisker tearing

Mice were lightly anesthetized (using 0.1 ml of the anesthesia mixture described above per mouse) and their whiskers were pulled off from both sides using tweezers and duct tape until all whiskers were removed.

##### Hearing loss

Mice (5 week-old) received a daily i.p. injection of 2 ml/kg BW gentamicin (50 mg/ml gentamicin sulphate, Biological Industries) for one week ^38,39^. Tests were performed 1-2 weeks after the injection period. Control mice received saline injections in the same manner.

##### Anosmia

Mice (8 week-old) received a single i.p. injection of 10 ml/kg BW methimazole (MMZ, Sigma-Aldrich) dissolved (10mg/ml) in sterile double-distilled water ^40,72^. Mice were subjected to all tests before receiving MMZ (control), and 1 week after MMZ treatment.

### Tissue preparation and immunostaining

#### Fixating and sectioning

Mice were perfused following i.p. injection of a ketamine and medetomidine mix (overdose of a 0.8 ml anesthesia mixture per mouse). Twenty ml of saline were passed through the heart, followed by 20 ml of 4% paraformaldehyde (PFA). Both cochleae were removed and placed overnight in 4% PFA at 4°C. For dissection of the nasal cavity, the mandibula was removed and the rest of the skull was placed overnight in 4% PFA. The following day, the cochleae and skulls were added to tubes containing 0.5 M EDTA, pH 8 (Sigma-Aldrich) for decalcification (∼5 days for cochlea, ∼7 days for skull). After the bones were decalcified, the organ of Corti was extracted from each cochlea and placed in Peel-A-Way embedding mold (Sigma-Aldrich) with 4% agar, positioned with apex facing upwards. The decalcified skulls were placed in the Peel-A-Way embedding mold with 4% agar, positioned with nostril facing up. Coronal sections (100 µm-thick) were sliced using a vibrotome (Leica VT1200 S) under a magnifying binocular. For slicing the organ of Corti, the knife amplitude was 0.7 mm and the slicing speed was 0.02 mm/sec; for slicing the nasal cavity, the knife amplitude was 0.5 mm and the slicing speed was 0.01 mm/sec.

#### Immunostaining for olfactory marker protein (OMP)

Sections containing the vomeronasal organ (VNO), main olfactory epithelium (MOE) and the olfactory bulb were processed for immunostaining using the following protocol. After a 30 min incubation in 0.3% Triton X-100 in PBS (PBS-t), the sections were incubated in blocking mix containing 20% normal goat serum (NGS) in PBS-t for 2 h. Then, the sections were placed in primary antibody mix (2% NGS, mouse monoclonal αOMP antibodies (1:500; Santa Cruz) and PBS-t) overnight at 4°C. The next day, the sections were washed 3 times for 10 min each with PBS, and then incubated with Alexa 488-conjugated secondary antibodies in PBS (1:500; Abcam) for 2 h.

The sections were subsequently washed 3 times with PBS, incubated for 3 min in DAPI solution (1:2000 DAPI (20 mg/ml); Sigma-Aldrich) and then washed again 3 times with PBS. The sections were placed on a slide (25 × 75 x 1.0 mm, superfrost plus, Fisherbrand) and once completely dry, were covered with mounting medium (Vectashield Hardset) and a coverslip (24 × 60 mm, Menzel-Glaser).

#### Phalloidin staining

Sections from the organ of Corti were incubated for 1 h at room temperature in a mix solution containing 10% NGS, phalloidin conjugated with Alexa 488 (1:2000) and PBS. After incubation, the sections were washed 3 times with PBS for 10 min each and then counter-stained with DAPI in the same manner as described above.

#### Image analysis

All fluorescence images were acquired with a Nikon A1-Red Confocal Microscope using 10×, 20x, and 40x objectives. To measure the mean fluorescence intensity (MFI), regions of interest of the same size (230 µm x 166 µm for the organ of Corti; 379µm x 133µm for the MOE; 912µm x 757µm for the VNO) was reconstructed using ImageJ analysis software. Average mean gray values (mean±SEM, arbitrary units) were calculated from 3 random sections from each subject, subtracting background values for the same image and normalizing by dividing the values obtained by the average mean fluorescence intensity of the background. For quantification following gentamicin treatment, 3 mice served as controls and 5 mice were injected with gentamicin. For quantification of the MOE following MMZ treatment, 3 mice served as controls and 4 mice were injected with MMZ. For quantification of the VNO following MMZ treatment, 4 mice served as controls and 3 mice were injected with MMZ.

### Statistical analysis

All statistical tests were performed using SPSS v21.0 (IBM). A Kolmogorov-Smirnov test was used to confirm the normal distribution of dependent variables. A one-tailed paired t-test was used to compare different conditions or stimuli for the same group, and a one-tailed independent t-test was used to compare a single parameter between two distinct groups. For comparing multiple groups and parameters, a mixed analysis of variance (ANOVA) model was applied to the data. This model contained one random effect (ID), one within-effect, one between-effect and the interaction between them. For comparison within a group using multiple parameters, a two-way repeated measures ANOVA model was applied to the data. This model contained one random effect (ID), two within-effects, one between-effect and the interactions between them. All ANOVA tests were followed, if the interaction or main effect were significant, by a post-hoc Student’s t-test. Significance was set at 0.05. When the normal distribution of variables was rejected, an equivalent non-parametric test was performed on the variables. The Mann-Whitney U test was performed instead of the independent t-test for comparing two distinct groups, and the Kruskal-Wallis test was performed instead of one-way ANOVA to compare the effect of the independent variable on the dependent variable (with 3 levels). The Friedman test was performed instead of the ANOVA repeated measures test for comparing the effect of the independent variable (each minute of the test) on the dependent variable (major movements). The Wilcoxon signed rank test was performed as a post-hoc test for the Friedman test and instead of the paired t-test for comparing two paired groups.

All statistical results for the various experimental groups used by us, including repetitions of the experiments, are detailed in Supp. Table 1.

## Supplementary Figures

**Supplementary Figure 1.**
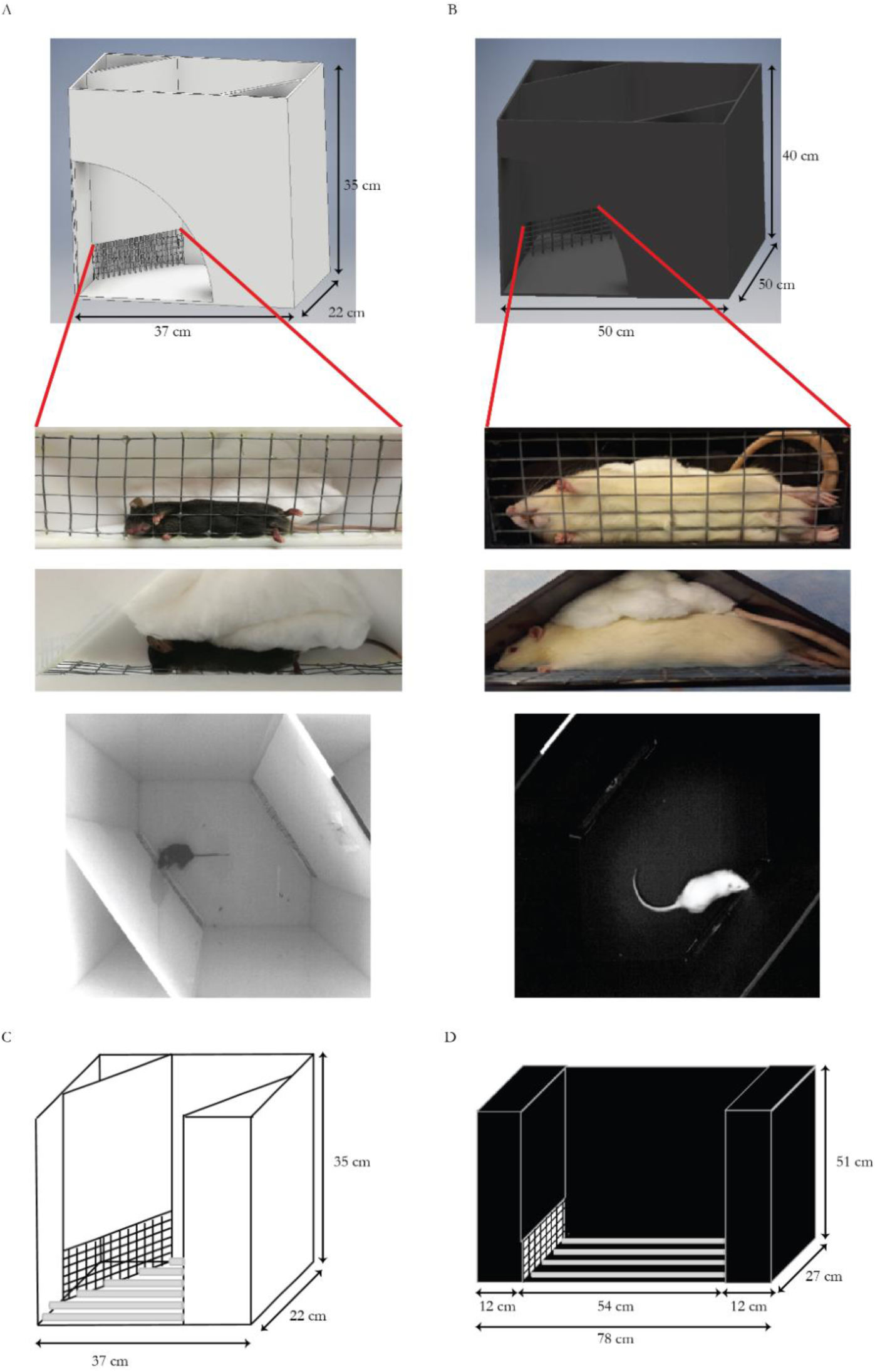
The behavioral systems used for social discrimination and social fear conditioning. (A-B) The behavioral arena, including triangular chambers for mice (left) and rats (right). The middle panels provide a closer view of the position of the anesthetized stimuli (front and upper views). The bottom panels show a frame from the video recording of the experimental setup during an SP test. (C) Scheme of the social fear conditioning arena used with mice. (D) As in C, for rats.

**Supplementary Figure 2.**
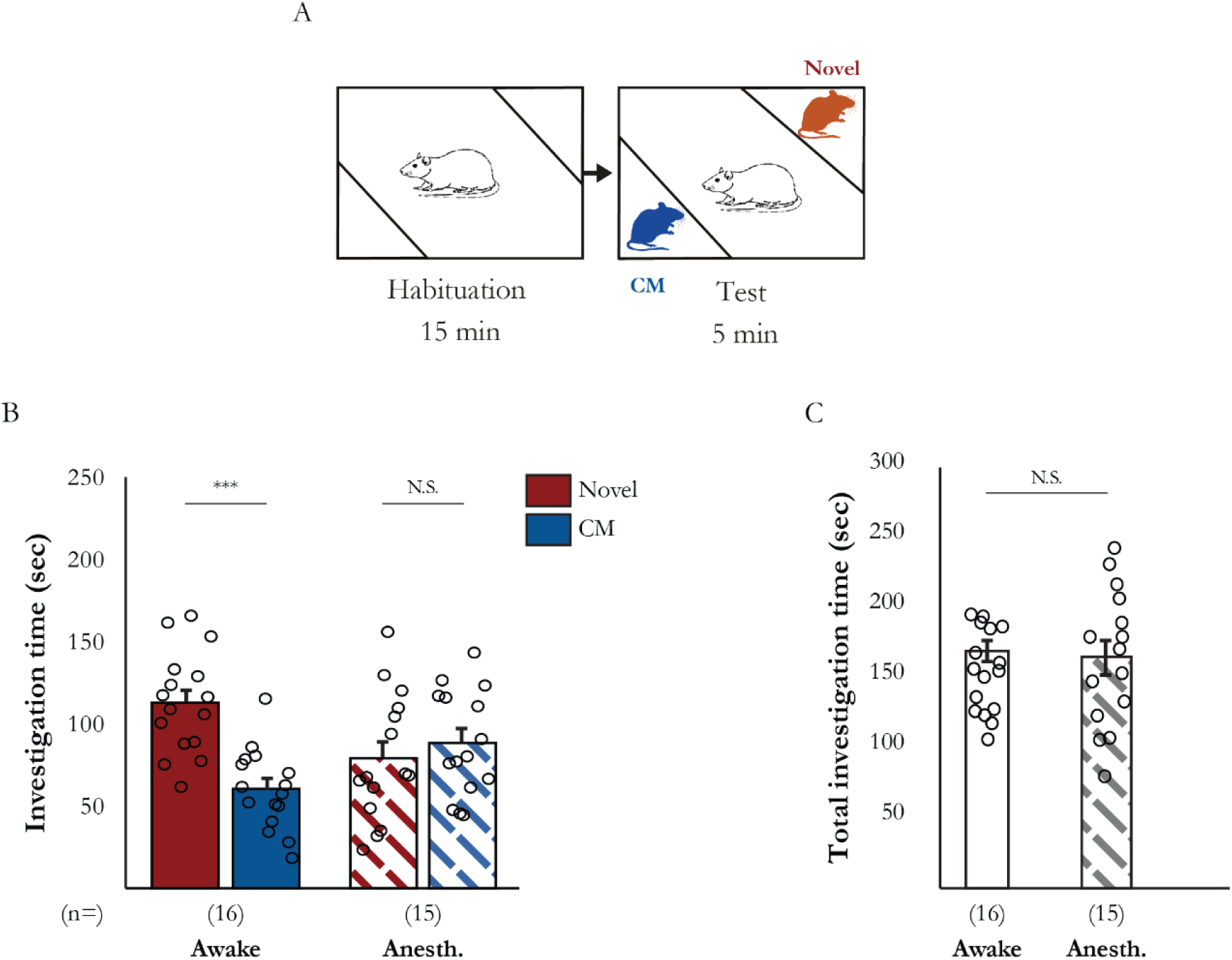
Social recognition of SD rats is impaired when anesthetized social stimuli are used. (A) Schematic representation of the familiarity discrimination paradigm with rats. (B) Mean investigation times for awake (filled bars) and anesthetized (dashed bars) stimuli in the familiarity discrimination test (awake: t(15)=4.536, p<0.001; anesthetized: t(14)=-0.716, p=0.243, paired t-test). (C) Mean total investigation time (of both stimuli) for awake (empty bar) and anesthetized (dashed bar) stimuli in the familiarity discrimination test (t(29)=0.336, p=0.370, independent t-test). ***p<0.001

**Supplementary Figure 3.**
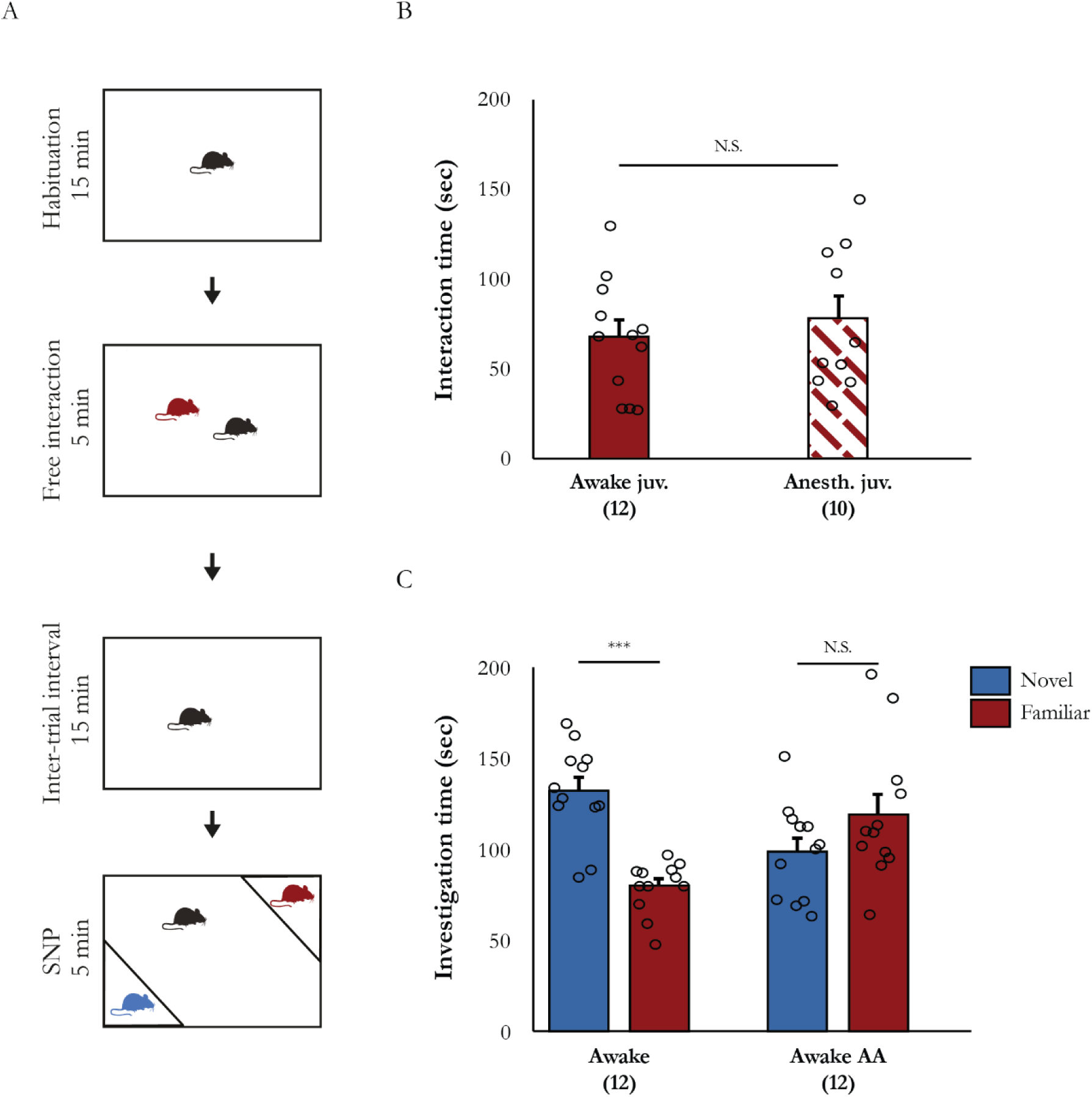
Adult male mice do not learn to recognize a social stimulus during free interaction while the stimulus is anesthetized. (A) Schematic description of the free interaction/SNP paradigm. (B) Mean interaction time of subjects with the juvenile stimulus during free interaction, using awake (filled bars) or anesthetized stimuli (dashed bars) (t(20)=-0.649, p=0.262, independent t-test). (C) Mean investigation time of juvenile stimuli during an SNP test, using such animals either awake or awake after anesthesia (Awake AA) (Stimulus x Group: F(1,22)=14.906, p<0.001, mixed model ANOVA. *post hoc* - awake: t(11)=5.534, p<0.001; anesthetized: t(11)=-1.263, p=0.117, paired t-test). ***p<0.001

**Supplementary Figure 4.**
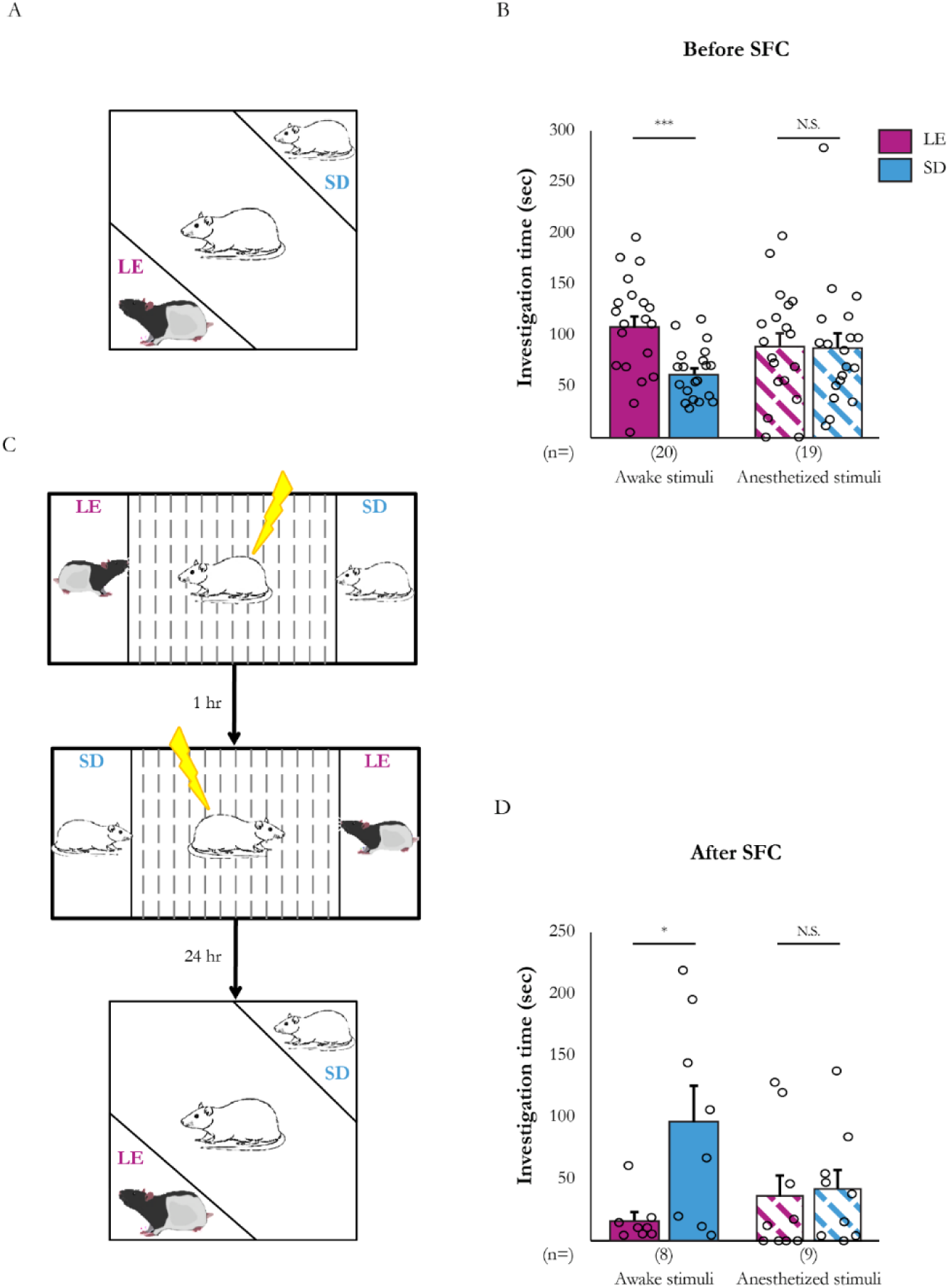
Impaired discrimination between anesthetized social stimuli following SFC. (A) Schematic description of the strain discrimination paradigm used with rats. Note that each stimulus is of a different strain. (B) Mean investigation time of both stimuli during the strain discrimination test before SFC, using awake (filled bars) or anesthetized stimuli (dashed bars) (Stimulus x Group: F(1,37)=3.268, p=0.040, mixed model ANOVA; *post hoc* - awake: t(19)=3.564, p=0.001; anesthetized: t(18)=0.063, p=0.475, paired t-test). (C) Schematic description of the SFC paradigm with rats. Note that we conducted two conditioning sessions with the two stimuli in opposite corners of the arena each time. (D) Same as in B, after SFC (Stimulus x Group: F(1,15)=3.749, p=0.036, mixed model ANOVA; *post hoc* - awake: t(7)=-2.780, p=0.014; anesthetized: t(8)=-0.266, p=0.399, paired t-test). *p<0.05, ***p<0.001

**Supplementary Figure 5.**
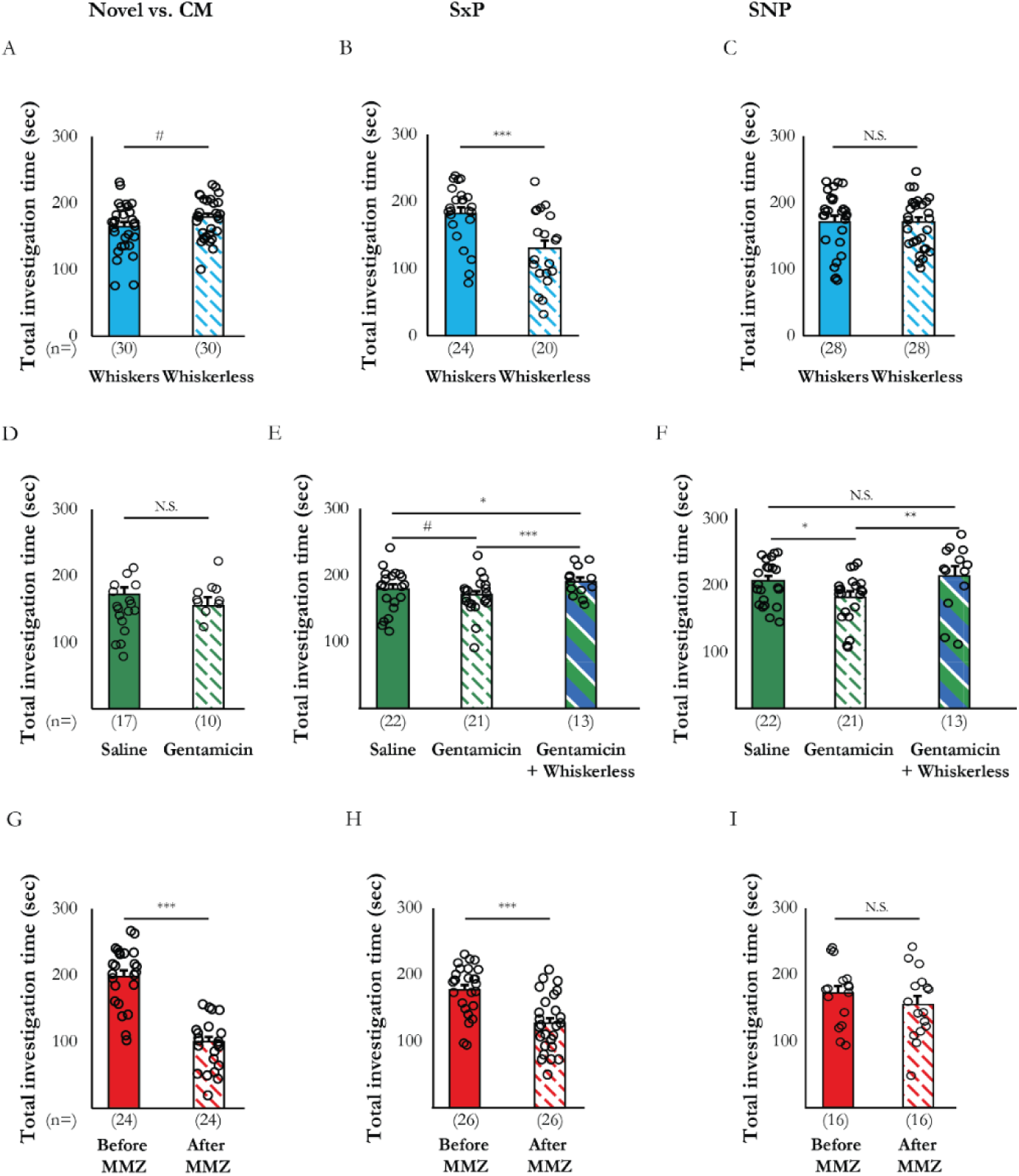
Total investigation time in the different tests after modality impairment. (A-C) Mean of total investigation time of subjects with (filled bars) and without (dashed bars) whiskers, in the familiarity discrimination test (A), the sex discrimination test (B) and the SNP test (C) (familiarity discrimination: t(29)=-1.689, p=0.051, paired t-test; sex discrimination: t(42)=3.853, p<0.001, independent t-test; SNP test: t(27)=0.336, p=0.370, paired t-test). (D-F) Mean of total investigation time of subjects injected with saline (filled bars), gentamicin (dashed bars) or of whiskerless mice injected with gentamicin in the familiarity discrimination test (D), the sex discrimination test (E) and the SNP test (familiarity test: t(25)=-1.307,p=0.101, independent t-test; sex discrimination: χ^2^(2)=12.151, p=0.001, Kruskal-Wallis test; *post hoc*: saline-gentamicin: U=171.5, p=0.074; saline-whiskerless+gentamicin: U=77, p=0.012; gentamicin-whiskerless+gentamicin: U=39, p<0.001, Mann-Whitney U test; SNP test: χ^2^(2)=7.088, p=0.015, Kruskal-Wallis test; *post hoc*: saline-gentamicin: U=158, p=0.038; saline-whiskerless+gentamicin: U=107, p=0.110; gentamicin-whiskerless+gentamicin: U=67, p=0.007, Mann-Whitney U test). (G-I) Same as in A, before (filled bars) and after (dashed bars) MMZ injection (familiarity test: t(46)=7.980, p<0.001, independent t-test; sex discrimination: t(25)=4.629, p<0.001, paired t-test; SNP test: t(15)=0.926, p=0.184, paired t-test). *p<0.05, **p<0.01, ***p<0.001

**Supplementary Figure 6.**
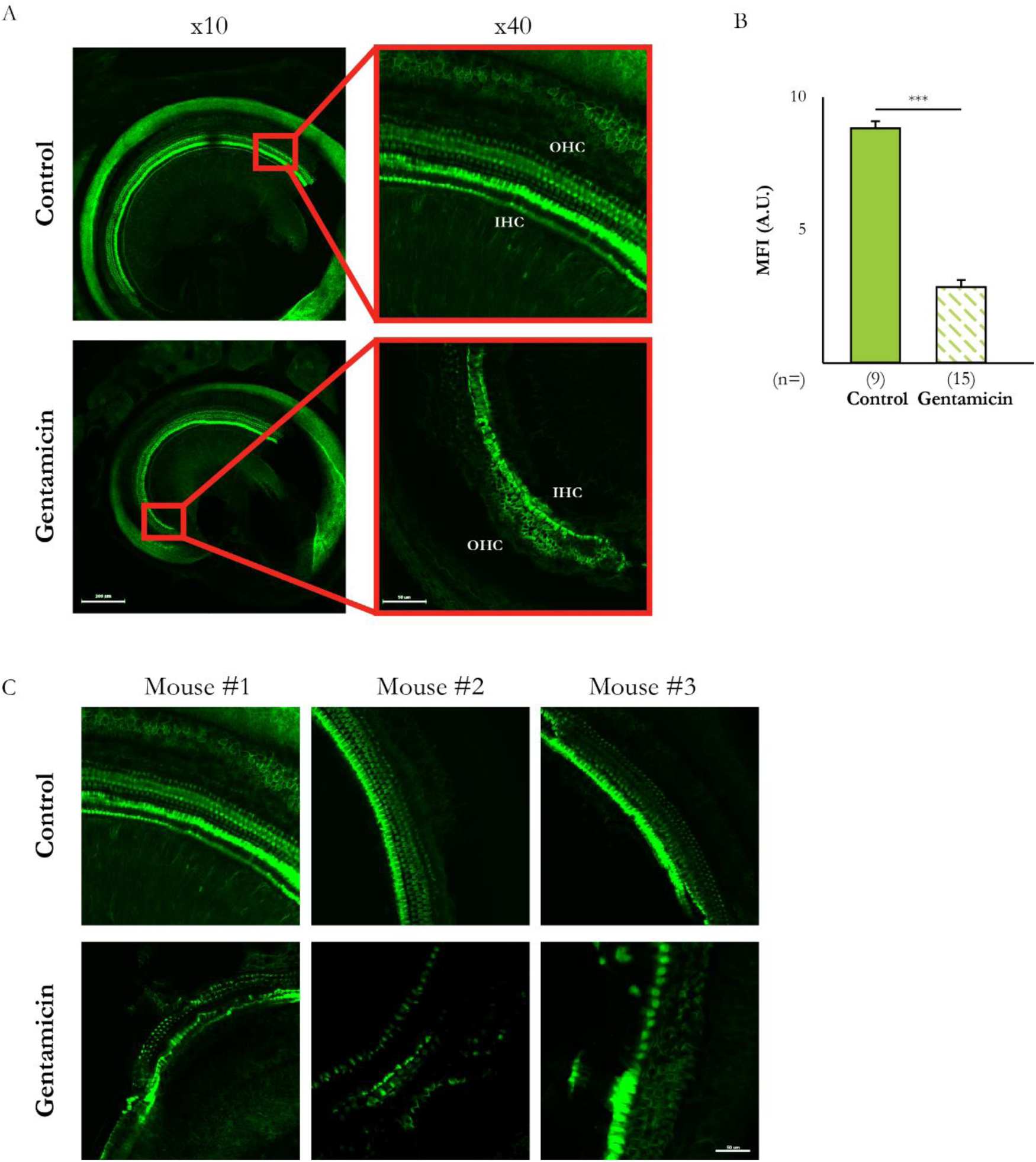
Cochlear hair cell destruction following gentamicin injection. (A) Representative images of the organ of Corti stained with phalloidin (green) from control (upper panels) and gentamicin-injected mice (lower panels), at low magnification (x10, left column, scale bar 200 µm) and higher magnification (x40, right column, scale bar 50 µm). Abbreviations: Outer hair cells (OHC), inner hair cells (IHC) (B) Mean fluorescence intensity (MFI) in arbitrary units (A.U.) in control (filled bars) and gentamicin-injected mice (dashed bars) (independent t-test, t(22)=13.005, p<0.001,). (C) Representative images from 6 different mice (3 control and 3 gentamicin-injected mice) allowing for comparison between the two groups at higher magnification (x40, scale bar = 50 µm). ***p<0.001

**Supplementary Figure 7.**
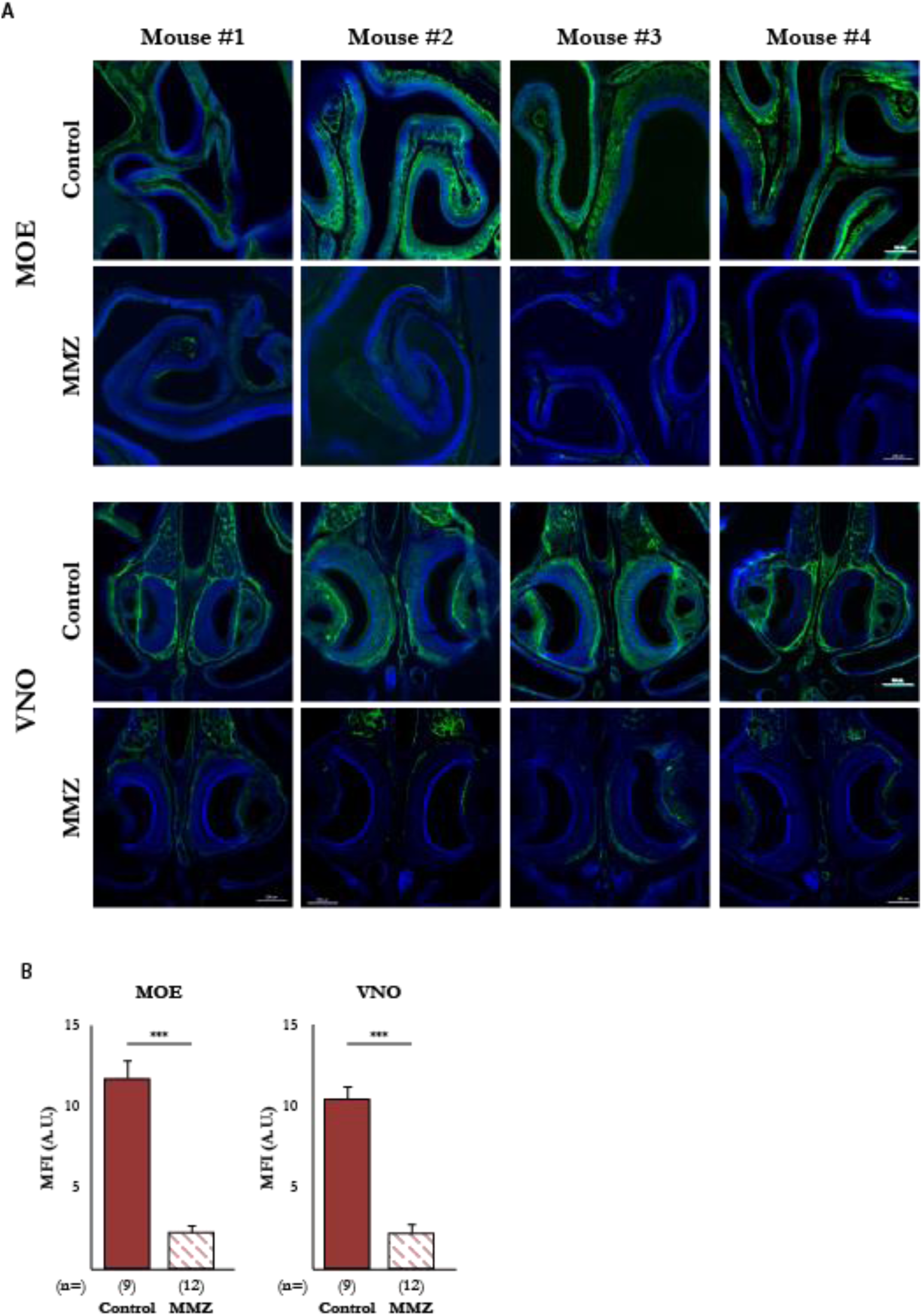
MMZ-mediated olfactory sensory neuron destruction in the MOE and VNO. (A) Representative images of the MOE (top set of panels) and VNO (bottom set of panels). OMP (green) and DAPI (blue) staining in the control group (upper panels) and MMZ-injected mice (lower panels), at low magnification (x10, scale bar = 200 µm). (B) Mean fluorescence intensity (MFI) in arbitrary units (A.U.) of control (filled bars) and MMZ-injected mice (dashed bars). (MOE: t(19)=8.647, p<0.001, independent t-test; VNO: t(19)=8.369, p<0.001, independent t-test). ***p<0.001

**Supplementary Figure 8.**
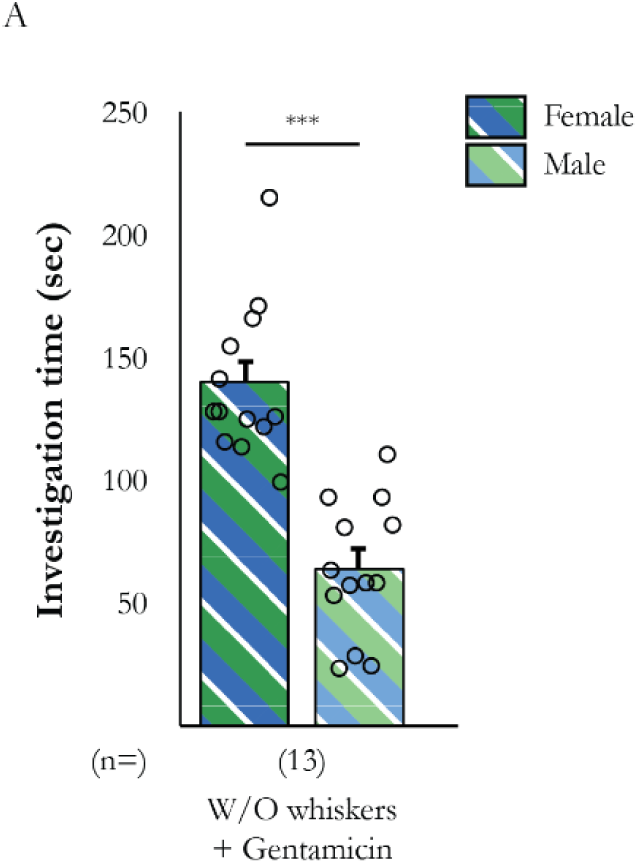
Sex discrimination is not affected by the combination of hearing and somatosensation loss. Mean investigation time of female (dark double-dashed line) and male (pale double-dashed line) stimuli by subjects in the sex discrimination test (t(12)=5.076, p<0.001, paired t-test). ***p<0.001

**Supplementary Figure 9.**
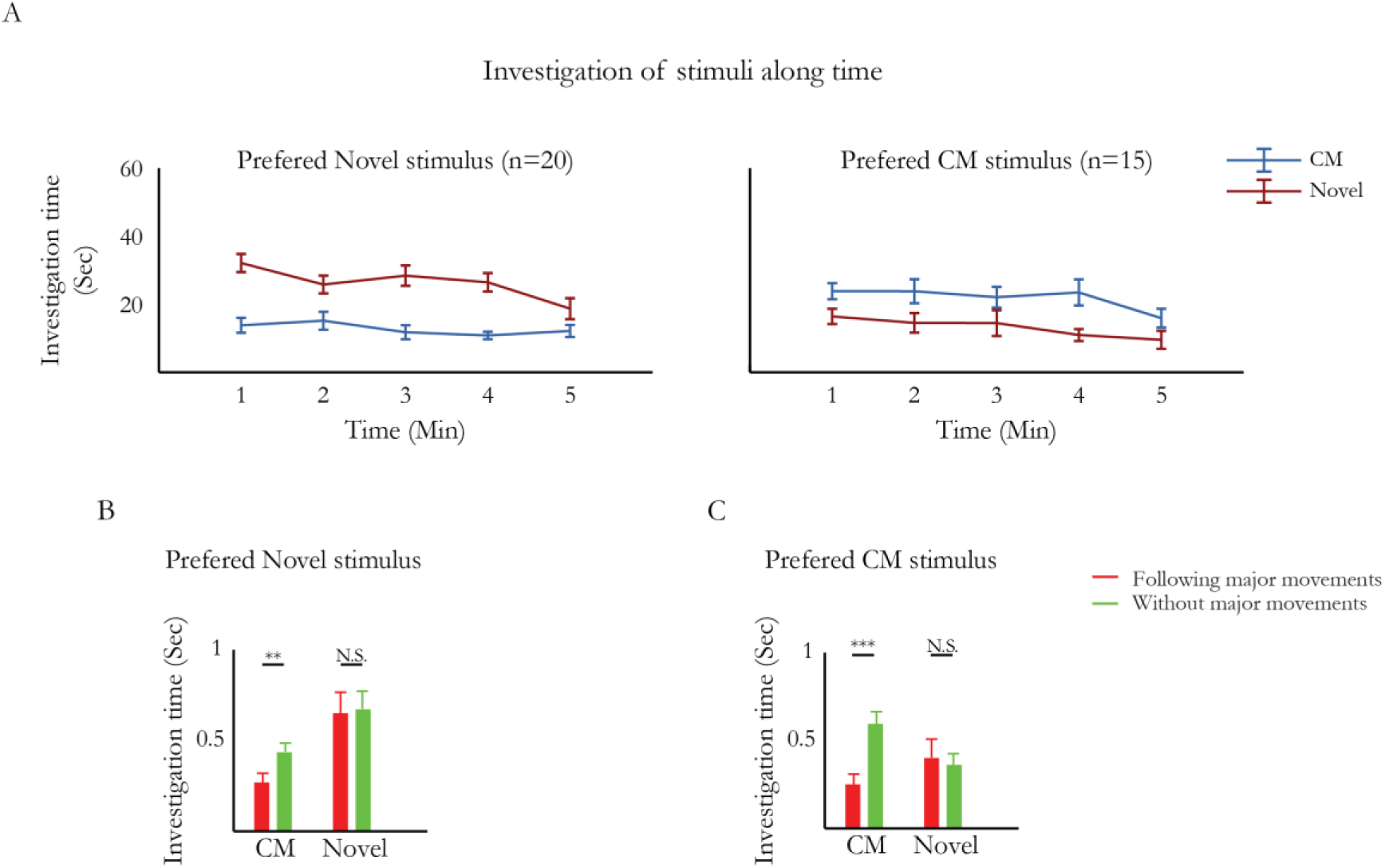
Comparison of response to movements of stimuli between subjects that preferred novel stimulus and those that preferred their CMs. (A) Mean investigation time of subjects in the familiarity discrimination test for a group of mice that preferred novel stimulus over their CM (left) and a group of mice that preferred their CM over novel stimulus (right), in 1 min bins. (B+C) Statistical analysis of the results shown in Figure 7I (interaction between the stimulus and the movements of the stimulus: F(1,29)=6.141, p=0.009, interaction between the groups, the stimulus and the movements of the stimulus: F(1,29)=1.166, p=0.144, 2 way -mixed ANOVA; *post hoc*-Preferred novel (B): CM movements: t(18)=-2.736, p=0.006, novel movements: t(17)=-0.194, p=0.424; Preferred CM (C): CM movements: t(12)=-4.088, p=0.001, novel movements: t(12)= 0.275, p=0.394, paired t-test). **p<0.01, ***p<0.001

**Supplementary Table 1.** Experimental groups.

| Paradigm | Condition | Repetitions | p value (t-test) |
| --- | --- | --- | --- |
| Familiarity test<br>(Related to Figure 1) | Both stimuli awake | Group 1 (n=20)<br>Group 2 (n=12) | p=0.001<br>p<0.001 |
|  | Both stimuli anesth. | Group 1 (n=20)<br>Group 2 (n=8) | p=0.151<br>p=0.467 |
| Sex preference test<br>(Related to Figure 1) | Both stimuli awake | Group 1 (n=12)<br>Group 2 (n=12) | p=0.036<br>p=0.006 |
|  | Both stimuli anesth. | Group 1 (n=18) | p<0.001 |
| Familiarity test<br>(Related to Figure 2) | Novel awake vs. CM anesth. | Group 1 (n=15) | p=0.189 |
|  | Novel anesth vs. CM awake | Group 1 (n=19)<br>Group 2 (n=20) | p=0.044<br>p=0.006 |
|  | CM anesth vs. CM awake | Group 1 (n=19) | p=0.002 |
|  | CM saline vs. CM awake | Group 1 (n=19) | p=0.205 |
| SP test<br>(Related to Figure 3) | Novel social stimuli awake | Group 1 (n=19)<br>Group 2 (n=20) | p=0.001<br>p=0.048 |
|  | Novel social stimuli anesth. | Group 1 (n=19)<br>Group 2 (n=19) | p<0.001<br>p<0.001 |
|  | CM social stimuli anesth. | Group 1 (n=16) | p<0.001 |
| SNP test<br>(Related to Figure 3) | Novel vs. familiar awake | Group 1 (n=19)<br>Group 2 (n=19) | p<0.001<br>p=0.010 |
|  | Novel vs. familiar anesth. | Group 1 (n=19) | p=0.225 |
|  | Novel vs. familiar AA | Group 1 (n=19) | p=0.467 |
|  | Novel vs. CM AA | Group 1 (n=16) | p=0.023 |
| Free interaction<br>(Related to supp. Figure 3) | Awake juv. stimulus | Group 1 (n=12) | p<0.001 |
|  | Anesth. juv. stimulus | Group 1 (n=12) | p<0.001 |
| SNP after free interaction<br>(Related to supp. Figure 3) | Both juv. stimuli awake | Group 1 (n=12) | p<0.001 |
|  | Both juv. stimuli AA | Group 1 (n=12) | p=0.116 |
| Before SFC<br>(Related to Figure 4) | C57bl/6j awake | Group 1 (n=8)<br>Group 2 (n=9) | p<0.001<br>p<0.001 |
|  | ICR awake | Group 1 (n=8)<br>Group 2 (n=9) | p=0.001<br>p<0.001 |
|  | C57bl/6j anesth. | Group 1 (n=10) | p=0.001 |
|  | ICR anesth. | Group 1 (n=10) | p=0.001 |
| After SFC<br>(Related to Figure 4) | C57bl/6j awake | Group 1 (n=8) | p=0.001 |
|  | ICR awake | Group 1 (n=8) | p=0.334 |
|  | C57bl/6j anesth. | Group 1 (n=9) | p=0.001 |
|  | ICR anesth. | Group 1 (n=9) | p<0.001 |
|  | C57bl/6j anesth. | Group 1 (n=10) | p=0.272 |
|  | ICR anesth. | Group 1 (n=10) | p=0.170 |
| Familiarity test<br>(Related to Figure 5) | Whiskerless | Group 1 (n=10)<br>Group 2 (n=20) | p=0.165<br>p=0.206 |
|  | Saline | Group 1 (n=7)<br>Group 2 (n=10) | p=0.031<br>p=0.020 |
|  | Gentamicin | Group 1 (n=10) | p=0.438 |
|  | After MMZ | Group 1 (n=10)<br>Group 2 (n=14) | p=0.218<br>p=0.329 |

**Supplementary Table 1 – Continuation**
| **Paradigm** | **Condition** | **Repetitions** | **p value (t-test)** |
| --- | --- | --- | --- |
| Sex preference test<br>(Related to Figure 5) | Whiskerless | Group 1 (n=20) | p<0.001 |
|  | Saline | Group 1 (n=12)<br>Group 2 (n=10) | p=0.004<br>p=0.015 |
|  | Gentamicin | Group 1 (n=11)<br>Group 2 (n=10) | p=0.008<br>p=0.013 |
|  | After MMZ | Group 1 (n=18)<br>Group 2 (n=8) | p=0.111<br>p=0.469 |
| SNP test<br>(Related to Figure 6) | Whiskerless | Group 1 (n=19)<br>Group 2 (n=9) | p=0.019<br>p=0.003 |
|  | Saline | Group 1 (n=12)<br>Group 2 (n=10) | p=0.003<br>p=0.062 |
|  | Gentamicin | Group 1 (n=11)<br>Group 2 (n=10) | p=0.180<br>p=0.001 |
|  | After MMZ | Group 1 (n=8)<br>Group 2 (n=8) | p=0.487<br>p=0.196 |

